# Beadex, the *Drosophila* LIM only protein, is required for the growth of the larval neuromuscular junction and sensorimotor activities

**DOI:** 10.1101/2023.09.20.558737

**Authors:** Kripa Chitre, Subhash Kairamkonda, Manish Kumar Dwivedi, Saumitra Yadav, Vimlesh Kumar, Sujit K. Sikdar, Upendra Nongthomba

## Abstract

The appropriate growth of the neurons, accurate organization of their synapses, and successful neurotransmission are indispensable for sensorimotor activities. These processes are highly dynamic and tightly regulated. Extensive genetic, molecular, physiological, and behavioural studies have identified many molecular players and investigated their roles in various neuromuscular processes. In this paper, we show that Beadex (Bx), the *Drosophila* LIM only (LMO) protein, is required for motor activities and neuromuscular growth of *Drosophila*. *Bx^7^*, a null allele, adult flies are flightless, with reduced walking and jumping activities. The larvae of *Bx^7^*, and the RNAi-mediated neuronal-specific knockdown of *Bx* show drastically reduced crawling behaviour, a diminished synaptic span of the neuromuscular junctions and an increased spontaneous neuronal firing with altered motor patterns in the central pattern generators (CPGs). Microarray studies identified multiple targets of Beadex that are involved in different cellular and molecular pathways, including those associated with the cytoskeleton and mitochondria, that could be responsible for the observed neuromuscular defects. With genetic interaction studies, we further show that *Highwire* (*Hiw*), a negative regulator of synaptic growth at the NMJs, negatively regulates *Bx*, as the latter’s deficiency was able to rescue the phenotype of the *Hiw* null mutant, *Hiw^DN^*. Thus, our data indicates that Beadex functions downstream of Hiw to regulate the larval synaptic growth and physiology.

## Introduction

A neuromuscular junction (NMJ) is a specialized chemical synapse that is formed between a presynaptic neuron and the post synaptic effector muscle. Synaptogenesis, the collective events that culminate in the formation of a synapse, is responsible for the appropriate assembly of the presynaptic components – active zones (AZs), neurotransmitter-filled synaptic vesicles (SVs), organelles like mitochondria, ion channels, etc. – along with the postsynaptic receptive field that chiefly includes the neurotransmitter-binding receptors, and their precise alignment with respect to each other. These processes are tightly governed by guidance molecules, morphogens as well as neuronal activity (Fox, 2009; Nakayama et al., 2005; K. Shen & Cowan, 2010; Williamson & Hiesinger, 2008). Impairment of any of these processes would have adverse effects on the NMJs that are often associated with clinical conditions like paralysis, Myasthenia gravis, and various motor neuron diseases. The lack of effective diagnostic and therapeutic approaches for such disorders prompts research in fundamental synapse biology.

*Drosophila melanogaster* is a well-studied model organism that not only offers a less complex neuromuscular system, but also draws striking parallels between the development, molecular regulators, and the synaptic transmission properties of chemical synapses to those of the vertebrates (Bellen et al., 2010). Studies in recent years have exploited the glutamatergic NMJs of *Drosophila* larvae, owing to their high similarity with glutamatergic synapses in the vertebral central nervous system (CNS), to study the roles of various genes whose mutations were subsequently associated with different human diseases. For example, Shaker and ether-a-go-go were found to be involved in the pathophysiology of cardiac arrhythmias, epilepsy and deafness (Jentsch, 2000); and synaptotagmin 2, which underlies Lambert-Eaton Myasthenic Syndrome was revealed to be a presynaptic Ca^2+^ sensor in fly studies (Herrmann et al., 2014). Further, various anterograde and retrograde signalling – Agrin, Jeb-Alk, Wnt, MAPK, Syt4, LAR-RPTP, Ephrin-Eph, TGF*β*, etc. – that were known to be involved in synapse development and growth were investigated by studying the morphological and/or physiological synaptic aberrations at the *Drosophila* larval NMJs, and behaviorally, by scoring these larvae for their characteristic abnormal crawling (Henríquez et al., 2011; Salinas, 2012; Salinas & Zou, 2008; Speese & Budnik, 2012).

One of the important classes of proteins working along this complex network of signalling is the transcription modulators. Multiple studies have associated them with the governance of synapse development, growth, and maintenance (Chédotal & Richards, 2010; Polleux et al., 2007; Ross et al., 2003). In the present study, we probed into the role of Beadex (Bx), the *Drosophila* LIM only (LMO) protein, whose mutant, *Bx^7^*, showcased strikingly reduced larval crawling activity, and flightlessness, reduced walking and jumping abilities in the adult flies. A neuron-specific deficiency of *Bx* resulted in a decreased synaptic span and size of synaptic boutons at the larval NMJs. These phenotypes were completely alleviated when genetically combined with *Hiw^DN^*as well as *Rae1^EX28^* mutations. This implies the interaction of *Bx* with *Hiw* and *Rae1* in regulating the NMJ morphology. Further, our electrophysiological experiments revealed stark aberrations in the spontaneous firing at the NMJs as well as the firing by the CPGs. In order to gain mechanistic insights, we performed a microarray analysis from the brains of the control and *Bx^7^* larvae. We found various cytoskeletal, ion channel, and mitochondria-associated genes to be differentially regulated that could hold plausible accountability towards the regulation of NMJs by *Bx*.

## Materials and Methods

### 1) Fly stocks and rearing

All fly stocks (Table S1) and crosses were maintained at 25 ± 2°C, unless mentioned otherwise, under a 12 h Light/Dark (L/D) cycle on standard cornmeal media (Lewis, 1960). The desired genetic combinations were attained by crossing virgin females with males, of the appropriate genotypes, in a 3:1 female to male ratio.

Crosses for all knockdown and overexpression experiments, were set and maintained at 18 ± 2°C; their progeny was harvested post-2 days of mating and shifted to 29 ± 2°C where they were incubated till the third instar stage. The same regime was observed for the mutants, as well, to maintain consistent conditions across experiments.

### 2) Larval crawling assay

The larval crawling assay was performed in a 100 mm petri plate containing 1% agar with a camera placed at a fixed height above it. Each larva was placed in the center of the petri plate and allowed to acclimate to its new surroundings, after which their movement was recorded for 1 min. The videos were converted to frames using Wondershare Filmora X and were further processed using ImageJ v1.8.0_172 (http://rsbweb.nih.gov/ij/). Subsequent analysis was performed using the wrMTrck plugin of ImageJ.

### 3) Larval fillet dissections

#### (i) For neuromuscular junction

Each wandering third instar larva was placed on a Sylgard plate with its dorsal side up and pinned, initially at its posterior end and then between the hooks at the anterior end, in cold HL3.1 buffer (70 mM NaCl, 5 mM KCl, 1.5 mM CaCl_2_, 5 mM MgCl_2_, 10 mM NaHCO_3_, 5 mM trehalose, 115 mM sucrose, and 5 mM HEPES [pH 7.2]) (Feng et al., 2004). A small horizontal incision was made, at the dorso-posterior side with a pair of fine scissors, which was then extended along the length of the larva from its posterior end to its anterior end (till the mouth hook). Two subsequent horizontal cuts were made on either side of this lengthwise cut at the anterior end. The larva was pinned, stretched, and spread (on all four sides) on the Slygard plate. The unwanted tissues were carefully removed with fine forceps, and the larval fillets were further processed for IHC.

#### (ii) For brain

Wandering third instar larvae of the appropriate genotype and gender were collected in a glass cavity block containing cold HL3.1 buffer. The larvae were pulled into two halves using a pair of fine forceps. The posterior half was discarded, and the anterior half was turned inside-out to expose the tissues inside. The brain was then teased free from the rest of the tissues and then further processed as required.

### 4) Immunohistochemistry and imaging

The dissected tissue was fixed using either of the following two methods, depending on their compatibility with the antigen and primary antibody involved – (i) 4% PFA in 1X PBS (Phosphate Buffer Saline) for 30 min or (ii) Bouin′s fixative for 5 min. The former fixation method was used for most of the IHC experiments, including staining with anti-CSP (mouse; 1:50; DSHB), anti-Repo (mouse; 1:10; DSHB), anti-pMAD (rabbit; 1:500; CST), and anti-Hiw (mouse; 1:50; DSHB) antibody, while the latter was explicitly used when the anti-GluRIIA (mouse; 1:50; DSHB) and anti-GluRIIC (rabbit; 1:50; A gift from Prof. DiAntonio) antibodies were used in the staining process; Alexa Fluor 488 conjugated anti-HRP antibody (that stains neuronal membranes; mouse; 1:500; Jackson Immunoresearch), and anti-BRP (mouse; 1:50; Jackson Immunoresearch) antibody were compatible with both methods. Post-fixation, the tissues were washed thrice with PBTx (0.3% Triton X-100 in 1X PBS) for approximately 15 min each. The tissues were subsequently blocked with 3% BSA (Bovine Serum Albumin) for 1 h at 25°C. The tissues were next, incubated with appropriate concentrations of the concerned primary antibodies overnight at 4°C. The unbound primary antibody was washed off using PBTx with mild rocking, three times for 10 min each. The tissues were further stained for the secondary antibodies – Alexa Fluor 568 anti-mouse (1:200), Alexa Fluor 594 anti-mouse (1:200), and Alexa Fluor 647 anti-rabbit (1:200) – washed, similarly. All secondary antibodies were sourced from Thermoscientific (Invitrogen). All antibody dilutions were made in 3% BSA. They were finally mounted on slides using VECTASHIELD mounting media. Images were captured using confocal microscopes – either LSM880 or Zeiss SP8. The NMJs, ventral nerve cord (VNC), and segmental nerves were captured at 63X magnification with a Z-stack interval of 0.5 μm, 3 μm, and 1 μm, respectively, whereas the whole brain was captured at a magnification of 40X with a Z-stack interval of 3 μm. Super-resolution imaging, for the GluRIIC-BRP juxtaposition and mitochondria structural studies, was performed on Leica Elyra7 and LSM980, respectively. All the images were processed and analyzed using ImageJ v1.8.0_172 (http://rsbweb.nih.gov/ij/).

### 5) Live Imaging

#### (i) Mitochondrial Imaging

Larvae expressing a GFP-fused mitochondrial membrane protein (mitoGFP) in a neuron-specific manner (*nSyb Gal4>UAS-mitoGFP*) were dissected as described in section 3 (i) on a Sylgard plate. Imaging of the mitochondrial movement (proxied by mitoGFP punctae) in segmental nerves innervating the A2 and A3 segments was performed using LSM 880 upright microscope, at a frequency of 0.5 Hz. The movement of the mitochondria was tracked using the MTrackJ plugin in ImageJ v1.8.0_172 (http://rsbweb.nih.gov/ij/).

#### (ii) Calcium Imaging

Brains from wandering third instar larvae expressing CaMPARI pan-neuronally (*nSyb Gal4>UAS-CaMPARI*), a fluorescent calcium sensor, were dissected as explained in section **Error! Reference source not found.** (ii) in a 35 mm glass-bottom dish (Cat. no. 81158; Ibidi GmbH, Germany) (Fosque et al., 2015). The dissection buffer (HL3.1) was then replaced with 0.2% low-melting agarose to help restrict the movement of the tissue while imaging. The setup was then bathed in cold HL3.1 buffer, as explained in Jayakumar et al., 2016. Images were obtained on an Epi-fluorescence Olympus DSU upright microscope as described previously (Fosque et al., 2015). Briefly, the brains were exposed to UV light (405 nm) for 5 s to photoconvert the fluorophore into its active form, and a time-lapse imaging was performed for a continuous stretch of 5 min at an interval of 5 ms each.

### 6) Electrophysiology

Electrophysiological recordings, for measuring the mEJPS and EJPs, were taken from muscle 6 of hemisegments, A2 or A3, as detailed in Imlach & McCabe, 2009. The wandering third instar larvae were dissected on a Sylgard plate in cold HL3.1 buffer without Ca^2+^ (dissection buffer) as described in section 3 (i). The brain was cautiously excised by holding it with a pair of forceps in one hand and snipping it free from the nerves with a pair of fine scissors in the other. The fine recording (borosilicate) electrodes with a resistance of 20-25 MΩ were pulled using a P-97 micropipette puller (Sutter Instruments) and back-filled with 3M KCl. For recordings, the dissection buffer was replaced with a cold HL3.1 buffer with 1.5 mM Ca^2+^ (recording buffer). Only muscles with resting membrane potentials ranging between -60 mV to -75 mV were selected for recordings. The mEJPs were recorded for 1 minute. For the EJPs, the inter-segmental nerve was stimulated, and the postsynaptic membrane potential was recorded for 1 min at a frequency of 1 Hz (Jan & Jan, 1976; Rikhy et al., 2002). To measure the input resistance, an ascending current was injected into muscle 6, using the recording electrode, from -300 pA to +300 pA with an increment step size of 50 pA. The corresponding variation in the membrane potential was noted, and an IV graph was plotted to obtain its input resistance.

To record the fictive locomotor activity of the larvae, the third instar larvae were dissected in the dissection media as mentioned in section 3 (i) with the modification of leaving the larval brain intact. Similar to the mEJP recordings, recording electrodes filled with 3M KCl were inserted in muscle M6 of the segments A2 or A3, and the changes in the postsynaptic potential were recorded for at least 30 min. During recordings, the dissection buffer was replaced with a cold HL3.1 buffer with 0.5 mM Ca^2+^. The Ca^2+^ levels were kept to a minimum to prevent strong contractions of the larva.

Data were digitized using a digital 1440 A and acquired with the help of pClamp 10 software (Axon Instruments, Molecular device, USA). AxoClamp 900A (Axon Instruments) was used to amplify the signal. The quantal content was calculated by dividing the average EJP amplitude with the average mEJP amplitude for each NMJ. The recordings were analyzed using Mini-Analysis Program 6.0 (Snapsoft) and WinWCP v5.6.2 software (Strathclyde).

### 7) Microarray studies, analysis and validation

A microarray was probed (Genotypic Technology) using RNA extracted from *w^1118^* and *Bx^7^* larval brains. The Trizol method was used, as per the manufacturer′s instructions (TRI Reagent, Sigma-Aldrich), for RNA extraction from three biological replicates. A *Drosophila melanogaster* whole-genome expression profiling chip of 4 X 44k format was used for probing the RNA. It possessed ∼3 probes for each transcript. The GeneSpring GX software, version 12.0, (Genotypic technology) and Microsoft Excel (MS Office 365), were used for data analysis. The data discussed in this publication have been deposited in NCBI’s Gene Expression Omnibus (Edgar et al., 2002) and are accessible through GEO Series accession number GSE197058 (https://www.ncbi.nlm.nih.gov/geo/query/acc.cgi?acc=GSE197058).

To validate the microarray results, 1 μg of the submitted RNA samples were converted into cDNA using a first-strand cDNA kit following the manufacturer′s protocol (Cat. No. K1622, Thermo Fischer Scientific). Reverse Transcriptase PCR (RT-PCR) was performed using HOT FIREPol® EvaGreen® qPCR Supermix and gene-specific primers, with *rp49* as the housekeeping gene. Each reaction was set in triplicates. The qRT-PCR was run in a StepOnePlus™ Real-Time PCR system (ABI). All the primers used are listed in **Error! Reference source not found.** S2. The 2^-ΔΔCt^ m ethod was used to compare results across the experimental samples. The log2 fold changes obtained were plotted using GraphPad PRISM 8.0.

### 8) Statistical Analysis

All statistical analyses were performed using GraphPad PRISM 8.0. Two datasets were compared using an unpaired Student’s *t*-test, whereas, in experiments involving the comparison of more than two datasets, one-way ANOVA along with an appropriate post hoc test (Tukey’s, Dunnett’s, or Sidak′s) was performed. Significance is represented as: ns, not significant; *, *P*<0.05; **, *P*<0.01; ***, *P*<0.001; ****, *P*<0.0001.

## Results

### 1) *Bx* mutants exhibit defective crawling ability in the larval stage

Characterization of *Bx^7^*, an amorphic mutant of *Bx,* had revealed defects in the crawling behavior of the *Drosophila* larvae implying the role of *Bx* in larval locomotion (Kairamkonda, 2014). We hence investigated the crawling behaviour in different mutants of *Bx* – the amorphic mutant, *Bx^7^*, and hypermorphic mutants, *Bx^1^*and *Bx^J^*. *Bx^7^* has a deletion of 2 kb in the genomic region that includes *Bx-RB* specific exon 1, the constitutive exons 2 and 3, and 70 bp of the 5′ UTR (Kairamkonda & Nongthomba, 2014). Of the two hypermorphs, *Bx^1^* exhibits phenotypes of weaker severity as only one of the two negative regulatory sites (located in the 3′ UTR of the *Bx* transcript) is ablated, while *Bx^J^* has both these sites ablated and hence is a more severe mutant (Shoresh et al., 1998; Zeng et al., 1998). The crawling ability was found to be severely compromised in *Bx^7^* (2.708 ± 0.11 cm) larvae as compared to their wild-type control, *w^1118^* (4.218 ± 0.14 cm), while it was significantly enhanced in *Bx^J^* (6.21 ± 0.21) with respect to its control, *Canton-S* (4.737 ± 0.22 cm) (Fig. 1). However, the distance crawled by *Bx^1^*larvae (5.002 ± 0.19 cm) was comparable to that of the control, *Canton-S* (4.737 ± 0.22 cm).

**Figure 1:**
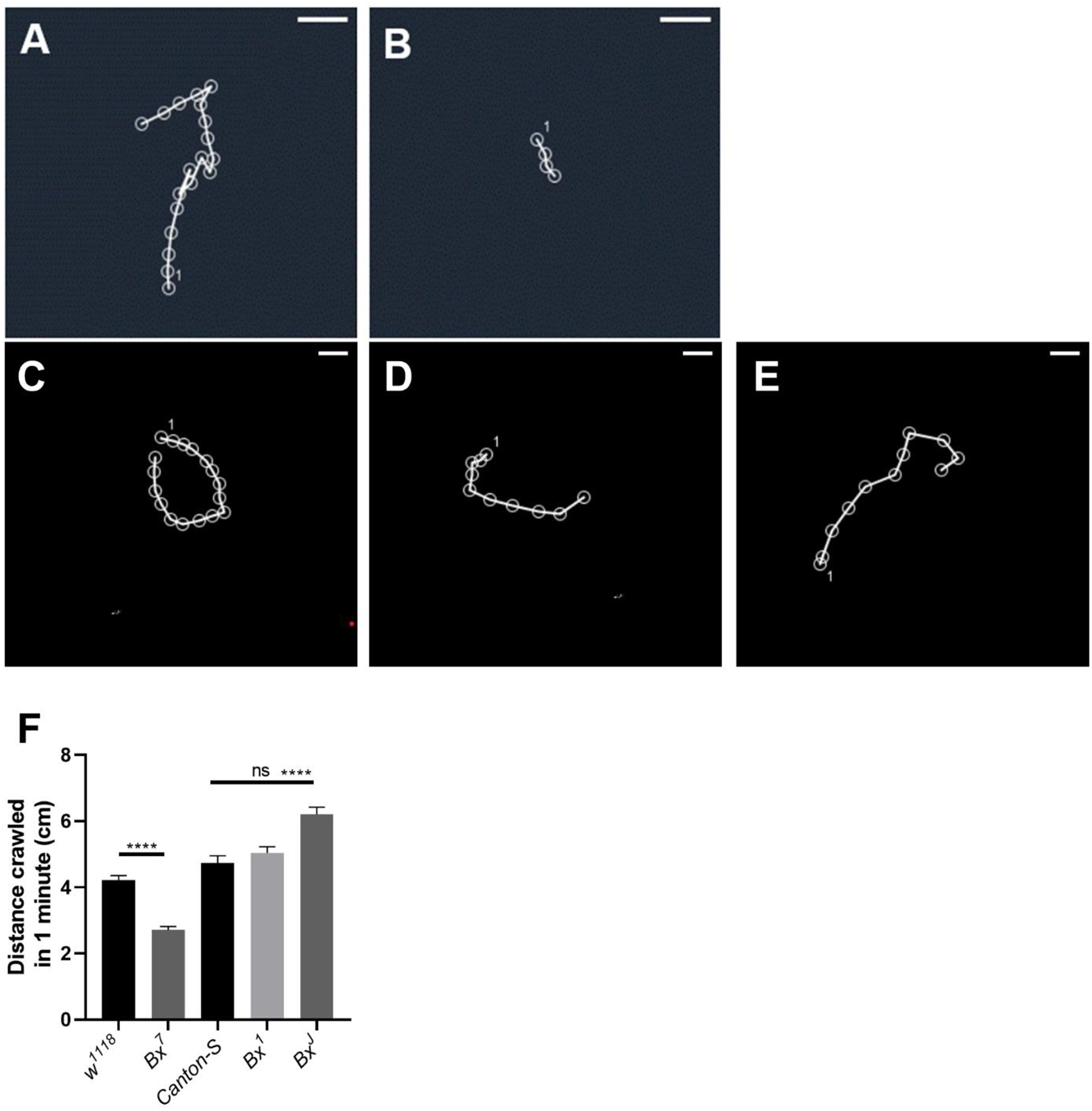
Bx mutation adversely affects the crawling ability of the larvae. Representative tracks of the path crawled by (A) control (*w^1118^*), (B) amorph (*Bx*^7^), (C) control (*Canton-S*), and hypermorphs (D) *Bx^1^*, and (E) *Bx^J^*. (F) Plot depicting the quantification of the distance crawled by these larvae. Scale bar represents 1 cm. For all the experiments, N=3, n>30. Statistical analysis was done using ordinary one-way ANOVA and a post hoc Tukey′s test. The error bar represents the standard error of the means (SEM), *** = p<0.001

### 2) *Bx* mutants have an abnormal synaptic span at the larval NMJs

Locomotor abnormalities are primary readouts for defective NMJs. To test our hypothesis, we imaged the NMJ structure at muscle 6/7 in segments A2 and A3 to extensively analyze various morphological features – NMJ span area, branch length, number of boutons, and number of branches – were quantified and compared. The anti-HRP (Horseradish Peroxidase) antibody was used to mark the neuronal membrane while presynapses were either labelled for CSP (Cysteine String Protein; a protein localizing in the SVs) or BRP (Bruchpilot; an active zone component) to individually identify each bouton. A significant reduction was observed in the NMJ span area (ratio of the area over which the presynaptic terminal was spread to the total area of the underlying muscles) in *Bx^7^* (0.19 ± 0.015) compared to that in *w^1118^* (0.26 ± 0.02) (Fig. 2A-A’). The NMJs of *Bx^7^* (30.73 ± 1.16) also report shorter branch lengths (length of the branch from the main branch to the terminal bouton) than controls (41 ± 1.55). The bouton numbers, bouton size and the number of branches did not vary significantly between *Bx^7^* and *w^1118^* (Fig. S2). Interestingly, the highest order of the branches was lower in the mutant (5^th^ order) vs the wildtype (6^th^ order), while their individual lengths remained unaltered (Fig. 2F).

**Figure 2:**
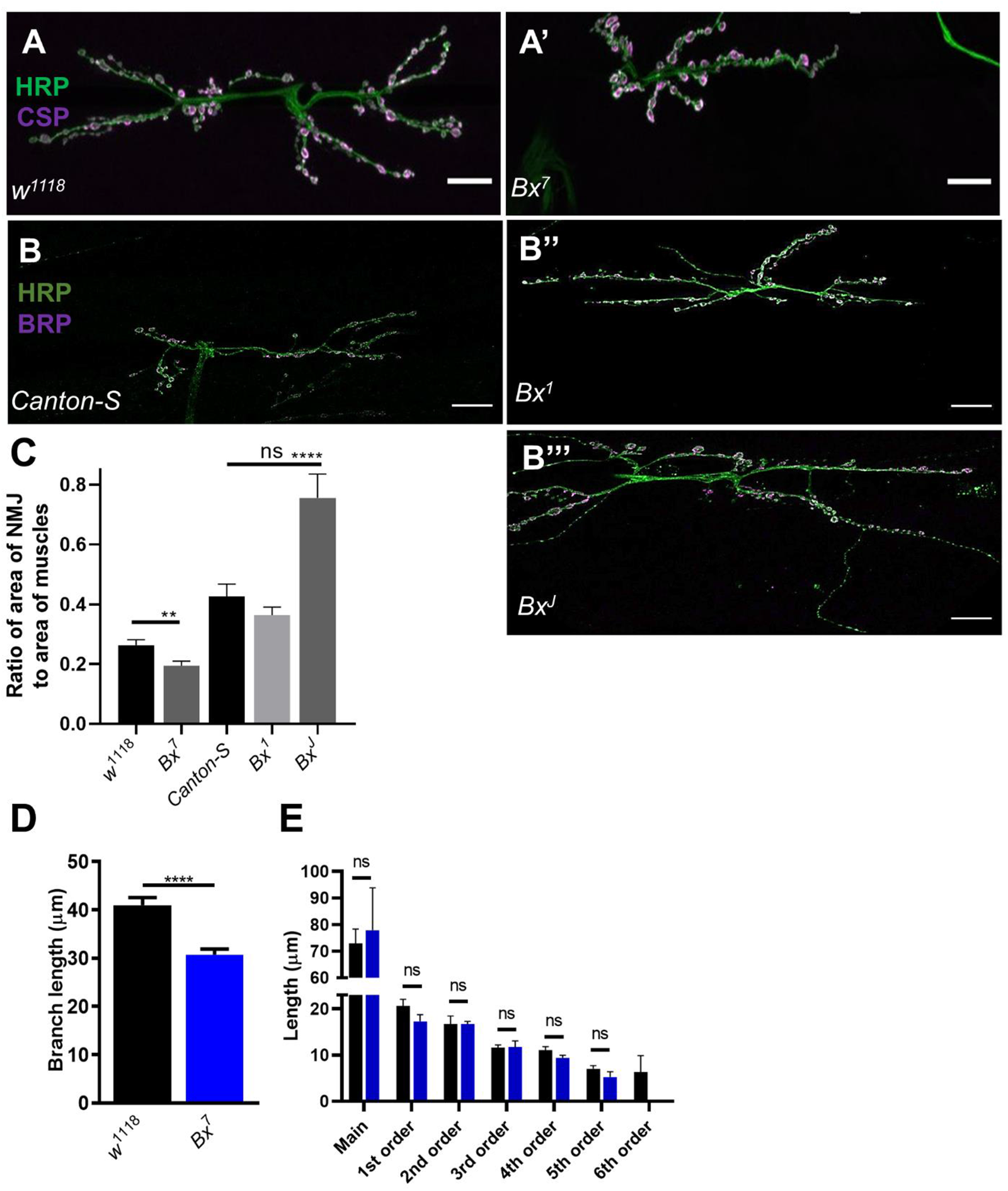
*Bx* mutant larvae show aberrations in branch length, synaptic span, and branch order. (i) Representative images of the NMJ at muscle 6/7 in (A) control (*w^1118^*) larvae vs (A’) in the amorph (*Bx^7^*). A significant reduction in (C) the NMJ span area and (D) the terminal branch length was reported. (E) Plot depicts the reduction in the highest order of branches with no variation in their individual lengths in *Bx^7^* compared to the controls. Green = anti-HRP, magenta = anti-BRP. Statistical analysis was done using the Student′s t-test. (ii) Representative images of the NMJ at muscle 6/7 in (B) control (*Canton-S*) larvae and the overexpression mutants (B’) *Bx^1^* and (B’’) *Bx^J^*. When compared to Canton-S, an increase in the (D) the NMJ span area was reported in the *Bx^J^* mutants. Green = anti-HRP, magenta = anti-CSP. Statistical analysis was done using ordinary one-way ANOVA and a post hoc Dunnett′s multiple comparison test. **** = p<0.0001, ns = not significant. Scale bar represents 20 μm. For all experiments, N=3, n>10. The error bar represents the standard error of the means (SEM), * = p<0.05, **** = p<0.0001, ns = not significant.

Similar to the behavioural phenotype, *Bx^J^* showed an opposite trend than *Bx^7^* when compared to its wild-type control; while *Bx^J^* mutant larvae (0.76 ± 0.08) had an expanded synaptic span area, the *Bx^1^* (0.36± 0.03) did not report any significant changes than *Canton-S* (0.43 ± 0.04) (Fig. 2B’-B’’’).

### 3) Bx is expressed in a subset of motor neurons

A defective NMJ may be an outcome of aberrations in the presynapse, or postsynapse, or both. In order to investigate which of these possibilities is probable, we studied the expression of *Bx* in the larval neuromuscular system. For this, we employed the well-established UAS – GAL4 genetic tool to label Bx expressing cells with GFP by driving *UAS-mCD8 GFP* (a construct that encodes for GFP localizing to the plasma membrane) using *Bx-Gal4*. The *Bx-Gal4* line had a P-element carrying a Gal4-regulatable enhancer and basal promoter inserted in the 2^nd^ intron of the *Bx* gene. We investigated its expression in the larval brain, in the presynaptic motor neuron, and the postsynaptic body wall muscle; the larval brain was counterstained with the glia-specific marker Repo and the motor neuron specific-marker pMAD; the presynaptic motor neuron was counterstained with anti-HRP; the postsynaptic body wall muscle was counterstained with phalloidin. This colocalization study revealed that *Bx* is expressed only in a subset of pMAD-expressing cells in the brain; it was completely absent in the glia, the motor neurons innervating the NMJs, and in the body wall muscles (Fig. 3). This suggested that Bx is expressed in the motor neurons and Bx in the neuronal component of the NMJs governs the NMJ morphology and the subsequent crawling of the *Drosophila* larvae.

**Figure 3:**
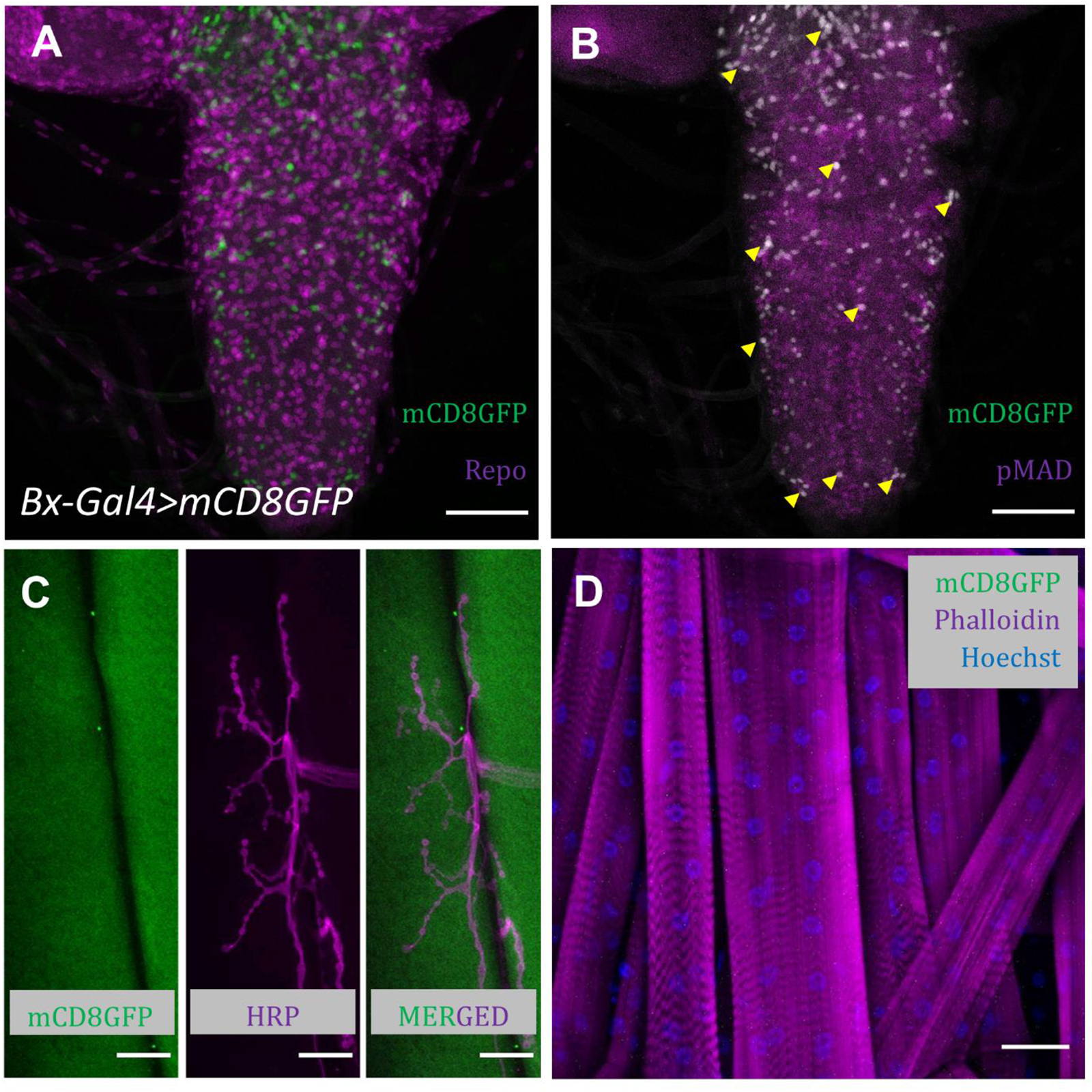
Bx is expressed in a subset of motor neurons. (A) *Bx-Gal4>UAS-mCD8GFP* (green) expression exclusive of repo (glial marker) expression (magenta). (B) Colocalization of *Bx-Gal4>UAS-mCD8GFP* (green) punctae with few pMAD (motor neuron marker) expressing neurons (magenta). The cells expressing both appear grey; a few of these cells have been marked with yellow arrows. Absence of GFP expression in (C) the presynaptic motor neuron at the NMJ and (D) the body wall muscles, when *UAS-mCD8GFP* expression was driven by *Bx-Gal4.* Scale bar represents 60 μm for (A) and (B) and 20 μm for (C) and (D).

### 4) Neuronal-specific knockdown of *Bx* recapitulates the behavioural and synaptic defects observed in *Bx^7^* larvae

In support of the above claim, we performed a tissue-specific knockdown of *Bx* by driving *UAS-Bx^RNAi^* using *Elav-GAL4; UAS-dcr2* (pan-neuron). The animals were scored for their crawling abilities and as anticipated, the mutant phenotype was recapitulated when *Bx* was knocked down in the neurons (Fig. 4A-C). This clearly suggested that *Bx* has a crucial role in *Drosophila* larval neurons. Since the neuronal knockdown was driven by *Elav-GAL4,* which is located on the X-chromosome, one would suspect variation in severity of the locomotor abnormalities between sexes due to effects of dosage compensation, and so male and female larvae were separately scored for their crawling ability, but surprisingly no significant difference was observed (Fig. S1D). Hence, data for males and females were pooled for all experiments henceforth.

**Figure 4:**
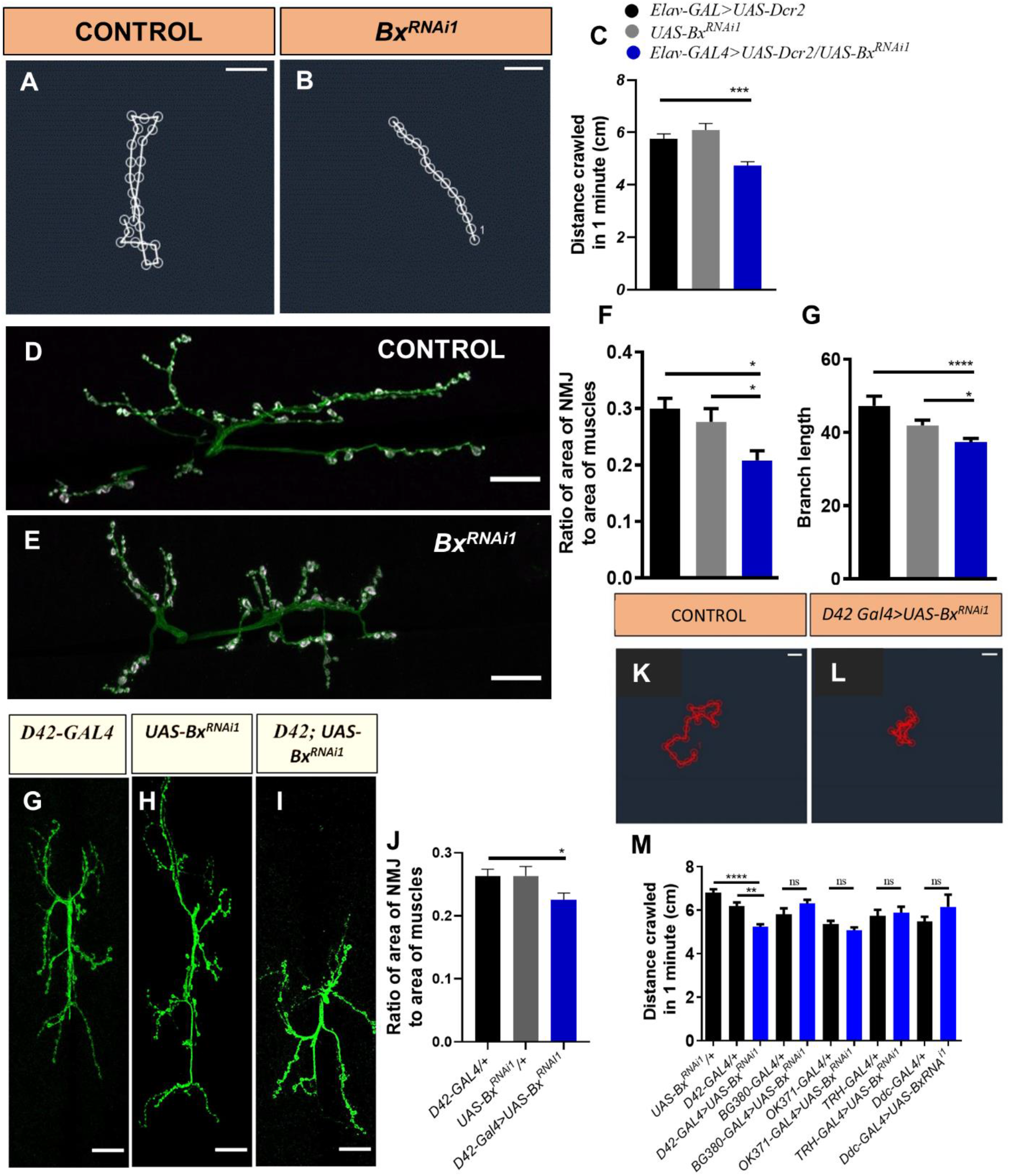
Neuronal-specific knockdown of *Bx* exhibits phenotypes in alignment with *Bx^7^*. Representative tracks of the path crawled by (A) control (*Elav-GAL4*) and (B) knockdown (*Bx^RNAi1^*). (C) Plot depicting the quantification of the distance crawled by these larvae. Scale bar represents 1 cm. For all the experiments, N=3, n>30. Representative images of the NMJ at muscle 6/7 in (D) control (*Elav-GAL4>UAS-Dcr2*) vs (E) in the knockdown (*Bx^RNAi1^*) larvae. A significant reduction in (F) the NMJ span area and (G) the branch length can be seen. Scale bar represents 20 μm. Green = anti-HRP, magenta = anti-CSP. For all experiments, N=3, n>10. Representative tracks of the path crawled by (K) control and (L) *D42-GAL4* driven knockdown of *Bx* larvae. (M) Comparison of population studies quantifying the distance crawled by larvae in which *Bx* knockdown was driven by *D42-GAL4, BG380-GAL4, OK371-GAL4, TRH-GAL4*, and *Ddc-GAL4*. (G-I) Representative images and (J) quantification show a reduction in the NMJ span area in larvae wherein *Bx* was knocked down by driving it using *D42-GAL4*. Scale bar in (K) and (L) represents 1 cm while that in (G-I) represents 20 μm. Green = anti-HRP. For the experiments, N=3, n>10. Statistical analysis was done using ordinary one-way ANOVA and a post hoc Tukey′s test. The error bar represents the standard error of the means (SEM), * = p<0.05, *** = p<0.001.

As *Bx* functions in the neurons and was expressed in a subset of motor neurons in the larval brain, we further aimed at identifying this subset. To achieve this, we knocked down *Bx* by driving *UAS-Bx^RNAi^*using five GAL4s that differed from each other in their spatio-temporal expression in the motor neurons. These motor neuron GAL4s include *D42-GAL4* (an insertion lying upstream of *toll-6* gene), *BG380-GAL4* (an insertion lying upstream of *futsch* gene), *OK371-GAL4* (an insertion lying in the proximity of *DVGLUT* gene), *TRH-GAL4* (inserted in the proximity of *tryptophan hydroxylase* gene), and *Ddc-GAL4* (inserted in the proximity of *dopa decarboxylase* gene) (Huser et al., 2012; Li et al., 2000; Mahr & Aberle, 2006; Sanyal, 2009). We scored these larvae for their behavioural phenotypes to assess which among them phenocopied *Bx* deficiency in larval neurons. A stark defect was observed in the crawling ability of the larvae when *Bx* was knocked down using *D42-GAL4* (*UAS-Bx^RNAi1^/+*: 6.81 ± 0.15; *D42-GAL4/+*: 6.19 ± 0.17; *D42-GAL4>UAS-Bx^RNAi1^*: 5.24 ± 0.11) (Fig. 4K-M). On the other hand, knockdown using all other GAL4s reported a crawling behaviour comparable to their respective controls (*UAS-Bx^RNAi1^/+*: 6.81 ± 0.15; *BG380-GAL4/+*: 5.81 ± 0.27; *BG380-GAL4>UAS-Bx^RNAi1^*: 6.31 ± 0.17; *OK371-GAL4/+*: 5.37 ± 0.14; *OK371-GAL4>UAS-Bx^RNAi1^*: 5.09 ± 0.13; *TRH-GAL4/+*: 5.75 ± 0.28; *TRH-GAL4>UAS-Bx^RNAi1^*: 5.89 ± 0.27; *Ddc-GAL4/+*: 5.48 ± 0.22; *Ddc-GAL4>UAS-Bx^RNAi1^*: 6.16 ± 0.57) (Fig. 4M). The behavioural phenotype associated with *D42-GAL4* driven knockdown was corroborated with the NMJ morphological phenotype, wherein a significant reduction was observed in the synaptic span compared to the controls (Fig. 4G-J).

### 5) *Bx* deficiency in the neurons causes an aberrant spontaneous firing at the NMJs and disrupts the rhythmic patterns generated by the CPGs

In addition to the structural defects at the NMJs, an aberrant crawling behaviour is often accounted for by defects in the synaptic transmission at these NMJs. In order to test this hypothesis, we performed electrophysiological studies at the NMJ of muscle 6/7 in segments A2 and A3. The postsynaptic potentials were recorded from muscle 6, and the following parameters were analyzed – mEJPs, EJPs, input resistance, and the rhythmic motor patterns. The mEJPs, EJPs, and input resistance were recorded from dissected fillets after excising the larval brain, while the motor patterns were recorded with intact brains. There was a stark increase in the frequency of mEJPs events in *Bx^7^* (frequency: 4.24 ± 0.21; amplitude: 0.87 ± 0.03) larvae compared to *w^1118^* (frequency: 2.40 ± 0.12; amplitude: 0.91 ± 0.04) while the neuronal knockdown of *Bx* (*Bx^RNAi^* larvae) resulted in a surge in both the frequency (*Elav-GAL4>UAS-dcr2*: 0.78 ± 0.06; *UAS-Bx^RNAi1^*: 0.69 ± 0.06; *Elav-GAL4>UAS-dcr2/ UAS-Bx^RNAi1^*: 1.12 ± 0.04) as well as the amplitude of the mEJPs (*Elav-GAL4>UAS-dcr2*: 0.79 ± 0.04; *UAS-Bx^RNAi1^*: 0.77 ± 0.04; *Elav-GAL4>UAS-dcr2/ UAS-Bx^RNAi1^*: 2.97 ± 0.18) than that in the controls (Fig. 5A, G-I). Intriguingly, the EJPs were not affected in either case of *Bx* deficiencies. Quantal content, the ratio of the average EJP amplitude to the average mEJP amplitude, was thence computed and was reported to be significantly reduced in the *Bx^RNAi^* larvae (*Elav-GAL4>UAS-dcr2*: 57.25 ± 1.96; *UAS-Bx^RNAi1^*: 63.04 ± 3.05; *Elav-GAL4>UAS-dcr2/ UAS-Bx^RNAi1^*: 37.06 ± 1.92) while it was unaltered in the *Bx^7^* larvae (*w^1118^*: 67.71 ± 1.77; *Bx^7^*: 61.40 ± 1.21), in comparison to their respective controls (Fig. 5B, C-E).

**Figure 5:**
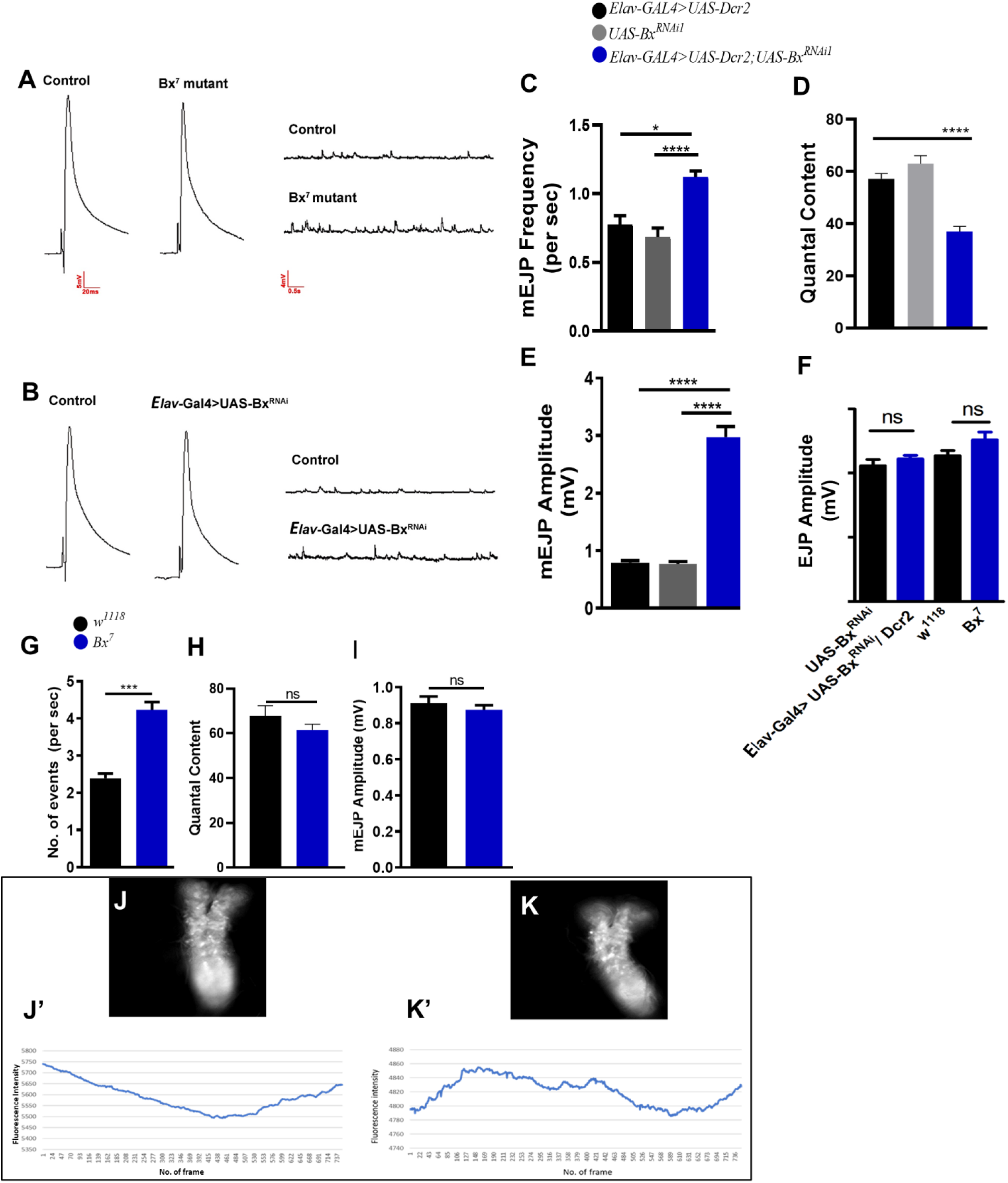
Increased spontaneous activity in *Bx* deficient larvae. (i) Representative traces comparing EJPs and mEJPs at muscle 6 of (A) *w^1118^* and *Bx^7^* larvae; (B) UAS-Bx^RNAi1^ and Bx^RNAi1^. Bar graphs represent the (C and G) mEJP frequency, (E and I) mEJP amplitude, and (D and H) quantal content, and the (F) EJP amplitude, as recorded from the postsynaptic muscle 6 at A2 and A3 hemisegments for the indicated genotypes. (ii) Representative grayscale images fluorescing red fluorescence post-activation by violet light for (J) *w^1118^* and (K) *Bx*^7^. The corresponding representative traces of calcium spikes with time are shown for (J’) *w^1118^*and (K’) *Bx*^7^ larval brains. Error bar represents the standard error of the mean (SEM). The statistical analysis was done using Student′s t-Test (for C-F) and one-way ANOVA followed by post-hoc Tukey′s test (for G-I). * = p<0.1, *** = p<0.001, **** = p<0.0001, ns = not significant.

The increase in the spontaneous activity may be attributed to three underlying causes: (i) aberrant transmission apparatus, (ii) reduced input resistance of the postsynaptic muscle, or (iii) aberrant motor patterns generated by the CPGs in the larval brain. We partially addressed the first hypothesis by immunostaining BRP and GluRIIA/GluRC to study the expression and localization of the presynaptic active zones and the postsynaptic glutamate receptor clusters, respectively. The number of BRP and GluRIIA/GluRC punctae per bouton were counted and no significant difference was observed in their numbers in the *Bx^7^* (BRP – *w^1118^*: 6.03 ± 0.13; *Bx^7^*: 7.2 ± 0.16, GluRIIA – *w^1118^*: 5.30 ± 0.14; *Bx^7^*: 5.77 ± 0.16, GluRC – *w^1118^*: 5.42 ± 0.12; *Bx^7^*: 5.72 ± 0.15) as well as the *Bx^RNAi^* larvae (BRP – *Elav-GAL4>UAS-dcr2*: 5.38 ± 0.25; *UAS-Bx^RNAi-1^*: 5.01 ± 0.16 *Elav-GAL4>UAS-dcr2/ UAS-Bx^RNAi-1^*: 5.20 ± 0.19, GluRIIA – *Elav-GAL4>UAS-dcr2*: 5.49 ± 0.12; *UAS-Bx^RNAi-1^*: 5.00 ± 0.16; *Elav-GAL4>UAS-dcr2/ UAS-Bx^RNAi-1^*: 4.98 ± 0.10, GluRC – *Elav-GAL4>UAS-dcr2*: 5.17 ± 0.15; *UAS-Bx^RNAi-1^*: 5.81 ± 0.27; *Elav-GAL4>UAS-dcr2/ UAS-Bx^RNAi-1^*: 5.29 ± 0.13) in comparison to their respective controls. Further, their localization, with respect to each other, did not differ upon *Bx* deficiency (Fig. S3). We next tested for the input resistance of the postsynaptic muscle by plotting its membrane potential changes to the input current steps. To avoid any systemic effect (esp. from the muscle), recordings were taken only for *Bx^RNAi^* larvae and its controls to reveal no significant differences (Fig. S3K).

Finally, we recorded the postsynaptic potentials in dissected fillets with intact brains to investigate the motor patterns generated by the CPGs for driving larval crawling. Striking variations were observed in motor patterns in *Bx^7^*vs those found in *w^1118^* larvae. Quantifications discovered significant reductions in the average interburst period (*w^1118^*: 14.12 ± 2.82; *Bx*^7^: 1.28 ± 0.07) and the burst amplitude (*w^1118^*: 5.16 ± 1.95; *Bx*^7^: 2.12 ± 0.21) in the mutants (Fig. 6). As all types of neurotransmissions – evoked, spontaneous, asynchronous – are dependent on intracellular Ca^2+^ spikes (Williams & Smith, 2018), aberrant spontaneous firing and motor patterns observed in *Bx* deficient larvae would hence have an underlying aberrant calcium spiking. To test our hypothesis, we performed live calcium imaging of larval brains expressing CaMPARI using *nSyb-Gal4*. Consistent with the electrophysiology results, calcium spikes in *Bx^7^*were found to have a higher frequency compared to the *w^1118^* (Fig. 5J-K’). These results further support the claim that *Bx* regulates the NMJ physiology, and the subsequent crawling, by regulating the motor patterns generated in the larval brain.

**Figure 6:**
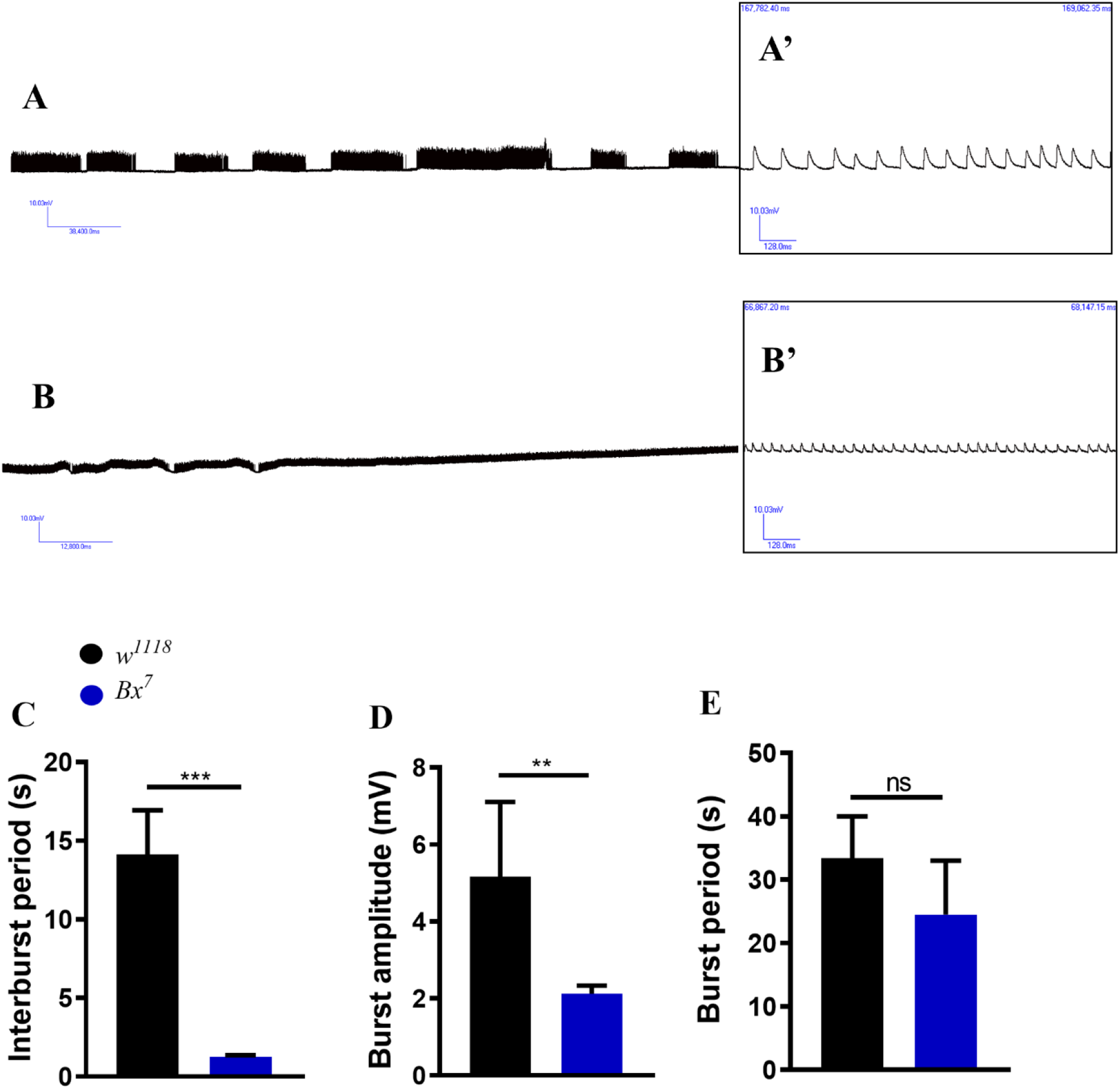
Aberrant motor patterns recorded from M6 of segment A2/ A3 during the fictive motion of *Bx^7^* larvae. Compressed representative traces of the bursts observed at muscle 6 of (A) *w^1118^* and (B) *Bx^7^* larvae. Zoomed-in traces during a burst are depicted in (A’) and (B’) for *w^1118^*and *Bx^7^,* respectively. Plots quantifying (C) interburst period, (D) Burst amplitude, and (E) Burst period. Error bar represents the standard error of the mean (SEM). The statistical analysis was done using Student′s t-Test (for C-F) and one-way ANOVA followed by post-hoc Tukey′s test (for G-I). ** = p<0.01, *** = p<0.001, **** = p<0.0001, ns = not significant.

### 6) *Highwire* negatively regulates *Bx* in the neurons of *Drosophila* larvae

Highwire (Hiw) is a ubiquitin ligase that has been reported to be a negative regulator of synaptic growth at the *Drosophila* larval NMJs (McCabe et al., 2004; Wan et al., 2000). *Hiw^DN^* mutants were found to have an increased NMJ span area (∼40%) as compared to the wild-type control larvae (∼18%) (Wu et al., 2005). The hypermorphic mutants, *Bx*^1^ and *Bx*^J^, exhibited a phenotype similar to that of *Hiw^DN^*, contrasting *Bx^7^*. This hinted toward a plausible negative interaction between the two genes in the larval neurons. To test the same, we employed a genetic approach wherein we knocked down *Bx* in the neurons of *Hiw^DN^* larvae (referred to as ‘Hiw rescue larvae’ hereafter) and compared their crawling ability as well as the NMJ morphology at muscle 6/7 with larvae of the following genetic makeup: (i) *w^1118^* (wildtype), (ii) *Hiw^DN^*, (iii) *UAS-Bx^RNAi-1^*/+ (wildtype), (iv) *Elav-GAL4>UAS-Dcr2* (wildtype) and (v) *Elav-GAL4*>*UAS-Bx^RNAi-1^*/*UAS-Dcr2*. Quantifications revealed that the average distance crawled in 1 min by the Hiw rescue larvae was restored to wildtype levels as opposed to the faster crawling observed in *Hiw^DN^* and the reduced crawling exhibited by the *Bx^RNAi^* larvae when compared to their respective controls (*w^1118^*: 3.80 ± 0.14, *Hiw^DN^*: 5.73 ± 0.23, *UAS-Bx^RNAi-1^*/+: 4.14 ± 0.22, *Elav-GAL4>UAS-Dcr2*: 4.36 ± 0.32, *Elav-GAL4*>*UAS-Bx^RNAi-1^*/*UAS-Dcr2*: 3.79 ± 0.26; *Elav-Gal4/Hiw^DN^; UAS-Bx^RNAi1^/UAS-dcr2*: 5.01 ± 0.21) (Fig. 7C). Further, the synaptic span at the NMJ in the Hiw rescue larvae was found to be comparable to that in wildtype, while *Hiw^DN^* had an expanded span and the *Bx^RNAi^* larvae had a reduced one (*w^1118^*: 0.22 ± 0.01, *Hiw^DN^*: 0.28 ± 0.01, *UAS-Bx^RNAi-1^*/+: 0.28 ± 0.02, *Elav-GAL4>UAS-Dcr2*: 0.29 ± 0.02, *Elav-GAL4*>*UAS-Bx^RNAi-1^*/*UAS-Dcr2*: 0.21 ± 0.01; *Elav-Gal4/Hiw^DN^; UAS-Bx^RNAi-1^/UAS-dcr2*: 0.25 ± 0.01) (Fig. 7A, D).

**Figure 7:**
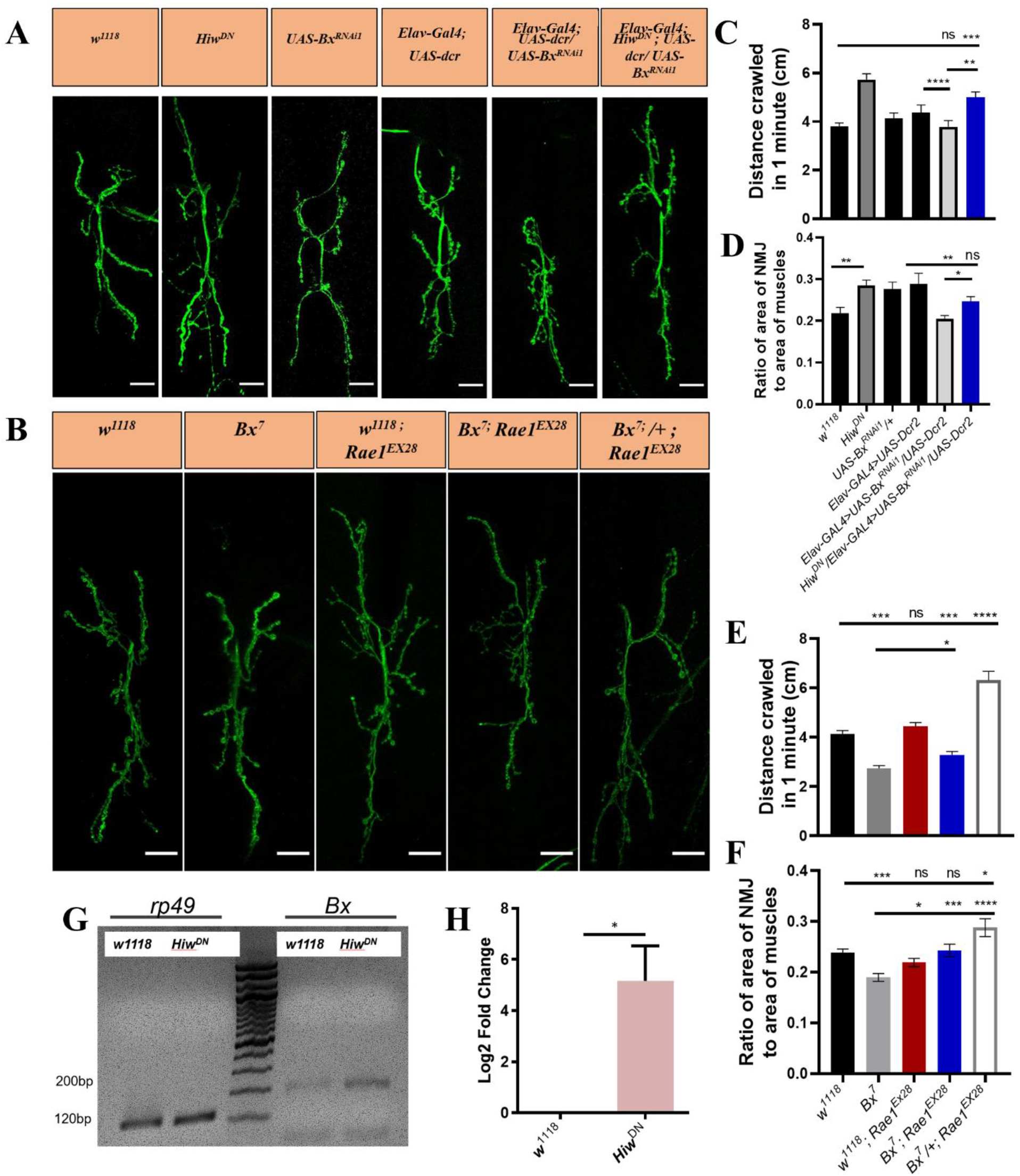
*Hiw^DN^* and *Rae1^EX28^* mutants rescue the *Bx^7^* phenotype. (i) Representative images of NMJs of muscles 6/7 of the following genotypes – (A) *w^1118^* (wild-type), *Hiw^DN^* (mutant), *UAS-Bx^RNAi1^*/+, *Elav-GAL4>UAS-Dcr2*, *Elav-GAL4*>*UAS-Bx^RNAi1^*/*UAS-Dcr2* (*Bx* knockdown), *Hiw^DN^/Elav-GAL4*>*UAS-Bx^RNAi1^*/*UAS-Dcr2* (*Bx* knockdown in *Hiw^DN^* mutant), (B) *w^1118^*(wildtype), *Bx*^7^ (mutant), *w^1118^*; *Rae1^EX28^*(mutant), *Bx*^7^; *Rae1^EX28^* (hemizygous *Bx^7^*with heterozygous *Rae1^EX28^*), *Bx*^7^/+; *Rae1^EX28^*(heterozygous *Bx*^7^ with heterozygous *Rae1^EX28^*). Plot depicts the quantification of the crawling assay performed to study the effect of (C) *Bx* knockdown in *Hiw^DN^* mutants and of (E) *Bx^7^* mutant allele in combination with the *Rae1^EX28^* allele. Plot depicts the quantification of the NMJ span area measured to study the effect of (D) *Bx* knockdown in *Hiw^DN^* mutants and of (F) *Bx*^7^ mutant allele in combination with the *Rae1^EX28^* allele. Scale bar represents 20 μm. Green = anti-HRP. For C & D experiments, N=3, n>30 while for E & F experiments, N=3, n>10. Error bar represents the standard error of the mean (SEM). Statistical analysis was done using ordinary one-way ANOVA and a post hoc Dunnett′s multiple comparison test for (C and D) while a post hoc Sidak′s multiple comparison test for (E and F). * = p<0.05, ** = p<0.01, *** = p<0.001, **** = p<0.0001, ns = not significant. (ii) (G) Representative agarose gel image exhibiting overexpression of *Bx* transcript levels in *Hiw^DN^* mutants compared to the controls (*w^1118^*). The constitutively expressed gene, *rp49*, was used as the internal control. (H) Plot showing the corresponding quantification of the upregulation of *Bx* transcript as revealed by the qPCR. For all experiments, N=3. Error bar represents the standard error of the mean (SEM). Statistical analysis was done using Student′s t-Test. * = p<0.05.

Further support was drawn by studying the genetic interactions between Rae1, a gene encoding a Hiw stabilizing protein, and *Bx.* We genetically combined *Bx^7^* and *Rae1^EX28^*, a hypomorphic mutant of *Rae1*, and analyzed their crawling as well as NMJ morphology. The hemizygous males (*Bx^7^* in its hemizygous state) and the heterozygous females (*Bx^7^* in its heterozygous state) larvae were scored separately. We found that the hyperactive crawling behaviour and the expanded synapse at the NMJ observed in the *Rae1^EX28^* mutants, and the reduced crawling and the decreased synaptic span observed in *Bx*^7^ were alleviated in the *Bx^7^*; *Rae1^EX28^* larvae (Fig. 7, E&F). Intriguingly, the *Bx^7^*/+; *Rae1^EX28^* heterozygous females outperformed not just the wildtypes but also the hemizygous *Bx*^7^; *Rae1^EX28^* larvae in both these assays (distance crawled in min – *w^1118^*: 4.14 ± 0.13, *Bx^7^*: 2.75 ± 0.10, *w^1118^*; *Rae1^EX28^*: 4.44 ± 0.16, *Bx^7^*; *Rae1^EX28^*: 3.30 ± 0.13, *Bx^7^*/+; *Rae1^EX28^*: 6.31 ± 0.36; synaptic span – *w^1118^*: 0.24 ± 0.01, *Bx^7^*: 0.19 ± 0.01, *w^1118^*; *Rae1^EX28^*: 0.22 ± 0.01, ^7^; *Rae1^EX28^*: 0.24 ± 0.01, *Bx^7^*/+; *Rae1^EX28^*: 0.29 ± 0.02).

To test the nature of the interaction between *Hiw* and *Bx*, we initially quantified the transcript levels of *Bx* in *Hiw^DN^*brains of the third instar larvae. The *Hiw^DN^* brains showed significantly higher levels of *Bx* transcripts as compared to that in *w^1118^* (Fig. 7G & H). Reciprocally, we analyzed the effect of Bx deficiency on the expression of *Hiw*. The quantification of Hiw protein levels, after immunostaining dissected third instar larval brains, found no significant variation between *w^1118^*and *Bx^7^* (Fig. S4).

### 7) *Bx* regulates the expression of the active zone-, ion channel- and mitochondria-associated genes in the larval neurons

*Bx* is a LIM-Only domain (LMO) containing protein whose role is that of a transcription co-activator or a scaffolding protein. This is suggestive of the fact that it would have a plethora of target genes. To identify them, we generated cDNA from larval brains of *w^1118^* and *Bx^7^* larvae and carried out a microarray study to find the differentially regulated genes in *Bx^7^* in comparison to *w^1118^*. A total of 420 genes were reported to be upregulated, while 676 genes were downregulated. To validate our results, we selected 7 highly downregulated genes and 6 highly upregulated genes and carried out a qPCR analysis and found that the results were in coherence with those of the microarray (Fig. 8B).

**Figure 8:**
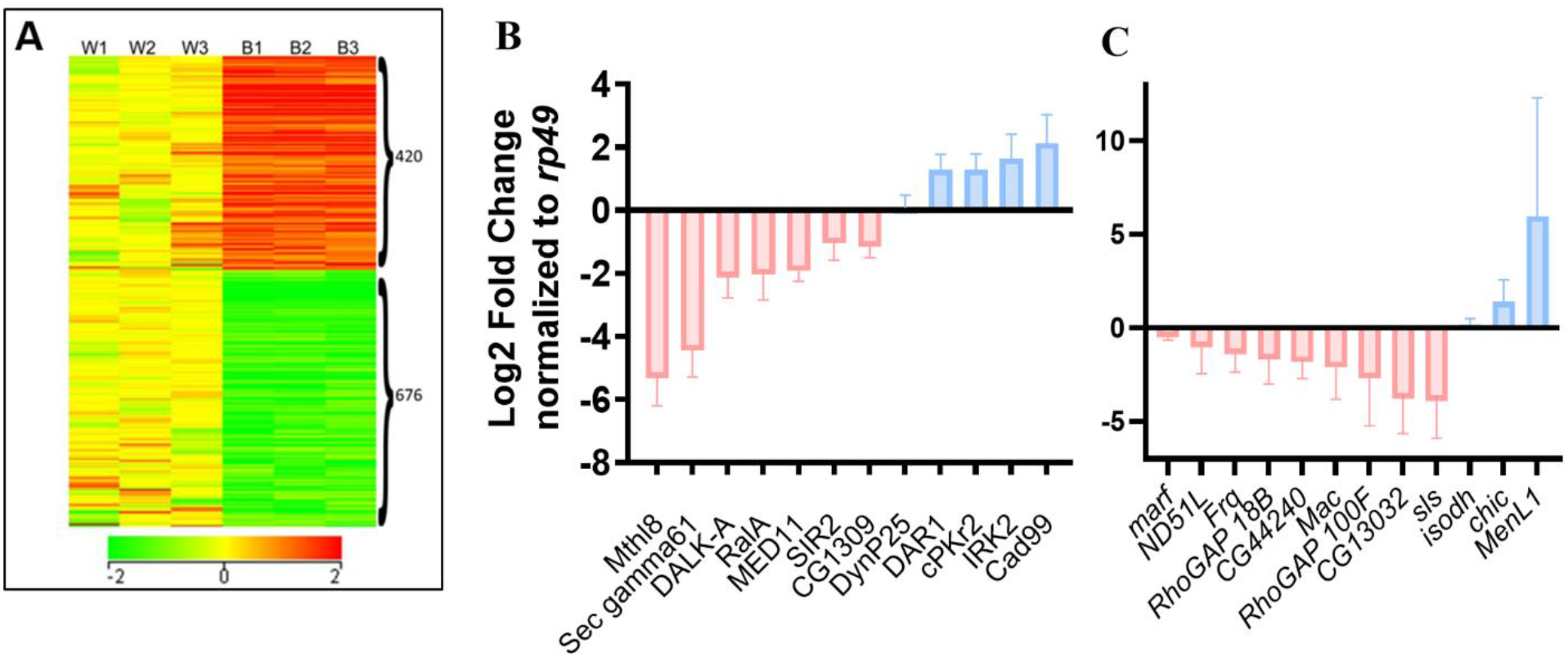
Microarray analysis of differentially regulated genes in the brains of *Bx^7^* larvae. (A) Heat map representation of the differentially regulated genes in larval brains of *Bx^7^* in comparison to *w^1118^*. (B) and (C) depict the qPCR quantification of the transcript expression levels of the selected candidate genes identified from the microarray study.

In order to identify downstream genes that would likely account for the morphological and physiological phenotypes observed in *Bx* mutant larvae, we further screened and clustered the genes into (i) actin-associated, (ii) microtubule-associated, (iii) mitochondria-associated, and (iv) ion-channel associated. A shortlist of 12 plausible candidate genes was validated using qPCR. All qPCR results were quantified using the ΔΔCt method while using *rp49* (*CG7939*) as the constitutive gene for normalization.

Among the differentially expressed genes that were associated with the cytoskeleton, the transcript levels of *Macoilin 1* (*CG30389*), a protein that is involved in both actin filament binding and microtubule-binding, were reduced ∼2 folds, that of *sls* (*CG1915*), a structural protein that enables actin-binding were reduced ∼3.8 folds, and that of *CG13032*, a putative microtubule organization protein, was found to have decreased by ∼3.7 folds. Three genes that are known to affect the actin polymerization were identified to be differentially regulated in the *Bx*^7^ larval brain – *RhoGAP 18B* (*CG42274*), a protein that regulates actin dynamics, was upregulated by ∼1.8 folds, *RhoGAP 100F* (*CG1976*), a protein known to regulate the cytoskeletal matrix organization was downregulated ∼2.7 folds, and *Chickadee* (*CG9553*), a protein that binds to actin monomers and is known to prevent actin polymerization was found to have an increased expression by ∼1.4 folds.

The next set of relevant genes that were found to be differentially regulated were the mitochondrial genes. Four such candidate genes were identified whose protein products were components of either the TCA cycle or the ETC or were involved in the fission/ fusion. *ND-51L* (*CG11423*; a predicted component of the mitochondrial respiratory complex I where it facilitates the transfer of electrons from NADH to ubiquinone) was reduced by ∼1.5 folds, and the *isocitrate dehydrogenase* (*NAD*(*+*)) (*CG5028*; an enzyme involved in the TCA cycle) transcript levels were decreased by ∼0.8 folds, *Malic enzyme-like 1* (*CG7964*; involved in malate metabolism) was found to be upregulated ∼6 folds, and *Marf* (*CG3869*; a GTPase belonging to the dynamin family that mediates mitochondrial fusion) was downregulated by ∼0.5 folds. *Marf* (Mitochondrial assembly regulatory factor) is involved in mitochondrial fusion, and its downregulation in the *Bx^7^* larval brains suggests the role of *Bx* in mitochondrial function and dynamics.

Lastly, genes associated with different ion channels in the larval neurons included the three most important genes, namely, *Irk2* (*CG4370*; an inward rectifying potassium channel) that was upregulated by ∼1.6 fold, *Frq1* (*CG5744*; a protein that modulates the synaptic efficacy in a Ca^2+^ dependent manner), and *CG44240* (a protein involved in divalent cation homeostasis) were found to be downregulated by ∼1.4 and ∼1.8, respectively.

*Marf* deficiency causes defective mitochondrial structure and distribution in neurons (Trevisan et al., 2018). With this lead, we looked at the mitochondrial distribution in *Bx^7^*larvae by expressing *UAS-mitoGFP* pan-neuronally, quantifying its intensity in the brain, and counting its puncta in the axons and the boutons. Intriguingly, we found that while the mitochondria were at comparable levels in the larval brains, their numbers were found to decline in the axons and increase significantly in the boutons of the NMJ compared to that of *w^1118^* (Fig. S5).

## Discussion

Bx, like other LMOs, is realized to have a diverse set of functions in a tissue-specific manner. In the current study, we have addressed the novel role of *Bx* in regulating the *Drosophila* larval NMJ structure and function.

We employed immunohistochemistry and genetics to study the expression pattern of Bx in the larval brain. Surprisingly, *Bx* did not express in the larval body wall muscles; unlike in adult IFMs (Mohan et al., 2015). Instead, Bx was found to be expressed in a subset of neurons expressing Toll-6, a member of the Toll-like receptor family involved in dendrite and axon guidance, and synaptic plasticity (Gaudet et al., 2011; Ulian-Benitez et al., 2017; Ward et al., 2015). We further showed that Bx deficiency resulted in aberrant motor patterns and NMJ structures in the larvae, suggesting that the Toll-6-expressing neurons may be involved in regulating the NMJ structures and motor output. However, the use of the UAS-GAL4 system in our experiments may have limitations in terms of spatiotemporal expression of GAL4 which may not imitate that of the protein under study. Therefore, further experiments such as fluorescence in situ hybridization (FISH) or immunohistochemistry staining could be performed to confirm our findings conclusively.

The deregulation of *Bx* exhibited striking defects in larval crawling ability. Any underlying muscular defect was ruled out as we demonstrated that Bx was not expressed in the larval body wall muscles (Fig. 3) and that neuron-specific knockdown of Bx phenocopied the mutant phenotype (Fig. 4). It is noteworthy that all the crawling assays were performed without distinctly stimulating the larvae, indicating an underlying defect in motor output rather than a sensory defect.

Our study reveals that Bx deficiency in neurons resulted in reduced synaptic span, characterized by decreased total branch length and branch order, but no significant difference in branch or bouton numbers. Bouton size, however, was affected only in the *Bx^RNAi^* larvae. Physiological investigations revealed a hyperactive spontaneous activity at the NMJs of *Bx* deficient larvae. Interestingly again, *Bx^RNAi^* larvae, unlike *Bx^7^* reported an increase in the amplitude of the mEJPs. The observed heightened severity in *Bx^RNAi^* as compared *Bx^7^* could have two underlying explanations – (i) the maternally deposited transcripts (if any) would be knocked out in case of the *Bx-UAS^RNAi1^*, as the knockdown is driven from the embryonic stage, while they shall persist in the mutant or (ii) the mutant would be facing a systemic level compensation.

Next, we investigated the molecular mechanisms underlying the role of Bx in NMJ regulation and found that Bx negatively interacted with Hiw, an E3 ubiquitin ligase enzyme that works as a part of the SCF (Skp/Cullin/F-box) complex to negatively regulate the MAP kinase signaling pathway through Wallenda (Wnd) (W. Shen & Ganetzky, 2009). The behavioural and morphological phenotypes resulting from Bx deficiency contrasted those observed in the *Hiw^DN^* and *Rae1^EX28^*larvae (Tian et al., 2011; Wan et al., 2000; Wu et al., 2005). However, molecularly, due to the lack of availability of an antibody against Bx, our study was limited to transcript levels.

The plausible downstream targets of Bx were suggested through the microarray experiment; it revealed the downregulation of RhoGAP100F (the *Drosophila* homolog of Syd-1), a critical AZ component that is essential for presynaptic development. It aids in the clustering of various presynaptic proteins like Neurexin, BRP, different scaffolding proteins, as well as mitochondria and synaptic vesicle docking (Hallam et al., 2002; Holbrook et al., 2012; Owald et al., 2010; Patel et al., 2006; Pilling et al., 2006; Wentzel et al., 2013). Previously, its deficiency was reported to reduce the NMJ span and expand the AZ (Owald et al., 2010). It can also be held responsible for the reduced quantal content due to reduced vesicle docking. We hypothesize that Bx deficiency may allow normal clustering of BRP, a presynaptic protein, despite the differential expression of RhoGAP100F, possibly because BRP requires the GTPase activity of RhoGAP100F to be recruited at the presynapse. However, we acknowledge that further experimental evidence is needed to support this hypothesis, such as ultrastructure imaging*. RhoGAP100F* and *RhoGAP18b* are known to function downstream of the BMP signalling pathway during the synapse regulation (Moustakas & Heldin, 2005). The BMP pathway works to positively regulate the synaptic growth; Gbb, Wit, and Tkv mutants develop small NMJs like *Bx* deficiency (McCabe et al., 2004; Rawson et al., 2003) while their upregulation effectuate an expanded synapse, like *Bx* hypermorphs (Dickman et al., 2006; Nahm et al., 2013; O’Connor-Giles et al., 2008). Moreover, Hiw plays a dual role and inhibits the BMP pathway to govern synaptic growth. In support, *LMO4*, a direct target of BMP7 and an interactor of MH1 and the R-Smads in the mammary epithelial cells, works along the BMP pathway in the vertebrate non-neuronal tissues (Wang et al., 2007; Lu et al., 2006). Hence, this indicates a plausible involvement of *Bx* in BMP signalling in the larval neurons.

IRK2 (Inwardly rectifying potassium channel 2) that hyperpolarizes the membrane was found to be upregulated in *Bx^7^*. It may hasten the restoration of the resting membrane potential allowing action potential firing at a faster rate. The upregulation of TCA cycle components isocitrate dehydrogenase (NAD(+)), & MenL1, and the ETC component ND-51L along with the increased mitochondrial density at the NMJs, would result in high amounts of ATP there. As ATPs are strongly associated with actin-dependent SV clustering, inhibition of Complex I, etc., this may lead to the hyperactive physiology found in *Bx^7^* (Lee & Peng, 2008). A biomolecular ATP assay would be more insightful of the same. Previously, *Bx* has been associated with circadian locomotor rhythms (Tsai et al., 2004). We, for the first time, report the involvement of Bx in the rhythmic motor patterns generated by the CPGs. Motor patterns in the *Drosophila* larvae were found to be plateau potentials when recorded in the form of postsynaptic muscle potentials (Kiehn & Kullander, 2004; Marder E. & Bucher D., 2001). These rhythmic bursts in specific circuitry help generate the inch-forward movement of the larvae. Since we could not identify which neurons were contributing to the aberrant calcium spikes, a direct cause and effect relationship between them would be difficult.

Lastly, the mitochondrion in the *Bx^7^* neurons showed tubular morphology in the soma, that could be associated with the downregulated transcript levels of Marf, a dynamin GTPase known to regulate mitochondrial fusion, and could affect its transport. Their shape and size, a result of fission and fusion events, dictates their quality as well as distribution. Furthermore, loss of Marf is known to affect synaptic transmission, develop morphological defects in the NMJs, and extend the larval lifespan supporting its contribution to other phenotypes observed in the *Bx*^7^ larvae (Sandoval et al., 2014). These results not only suggested the fact that *Bx* is involved in mitochondrial dynamics but also highlights its plausible involvement in the retrograde transportation of mitochondria in motor neurons.

## Conclusion

LMOs act as scaffolding proteins, to bring about a cell/tissue-context-specific gene expression (Milan et al., 1998; Zenvirt et al., 2008). Here, we have exploited the *Drosophila* larval model system, a convenient and strong genetic tool, to uncover the neuron-specific role of Beadex, the *Drosophila* LMO, with respect to the structure and function of the NMJs. We show plausible molecular pathways which could be important for Beadex phenotypes and neuromuscular development. Our results also indicate that Beadex may be involved with the Highwire signalling pathway.

## Conflict of Interest

Authors declare no conflict of interest.

## Notes

https://www.ncbi.nlm.nih.gov/geo/query/acc.cgi?acc=GSE197058

### Competing Interest Statement

The authors have declared no competing interest.

## References

Bellen, H. J., Tong, C., & Tsuda, H. (2010). 100 Years of Drosophila Research and Its Impact on Vetebrate Neuroscience. Neuroscience, 11(7), 514–522. 10.1038/nrn2839.100

Chédotal, A., & Richards, L. J. (2010). Wiring the brain: The biology of neuronal guidance. Cold Spring Harbor Perspectives in Biology, 2(6), 1–17. 10.1101/cshperspect.a001917

Dickman, D. K., Lu, Z., Meinertzhagen, I. A., & Schwarz, T. L. (2006). Altered synaptic development and active zone spacing in endocytosis mutants. Current Biology, 16(6), 591–598. 10.1016/j.cub.2006.02.058

*Drosophila media preparation | BangaloreFLY*. (2017). https://bangalorefly.ncbs.res.in/drosophila-media-preparation

Feng, Y., Ueda, A., & Wu, C. F. (2004). A modified minimal hemolymph-like solution, HL3.1, for physiological recordings at the neuromuscular junctions of normal and mutant Drosophila larvae. Journal of Neurogenetics, 18(2), 377–402. 10.1080/01677060490894522

Fosque, B. F., Sun, Y., Dana, H., Yang, C. T., Ohyama, T., Tadross, M. R., Patel, R., Zlatic, M., Kim, D. S., Ahrens, M. B., Jayaraman, V., Looger, L. L., & Schreiter, E. R. (2015). Labeling of active neural circuits in vivo with designed calcium integrators. Science, 347(6223), 755–760. 10.1126/science.1260922

Fox, M. A. (2009). Development of the vertebrate neuromuscular junction. The Sticky Synapse: Cell Adhesion Molecules and Their Role in Synapse Formation and Maintenance, 39–84. 10.1007/978-0-387-92708-4_3

Gaudet, P., Livstone, M. S., Lewis, S. E., & Thomas, P. D. (2011). Phylogenetic-based propagation of functional annotations within the Gene Ontology consortium. Briefings in Bioinformatics, 12(5), 449–462. 10.1093/bib/bbr042

Hallam, S. J., Goncharov, A., McEwen, J., Baran, R., & Jin, Y. (2002). SYD-1, a presynaptic protein with PDZ, C2 and rhoGAP-like domains, specifies axon identity in C. elegans. Nature Neuroscience, 5(11), 1137–1146. 10.1038/NN959

Henríquez, J. P., Krull, C. E., & Osses, N. (2011). The Wnt and BMP families of signaling morphogens at the vertebrate neuromuscular junction. International Journal of Molecular Sciences, 12(12), 8924–8946. 10.3390/ijms12128924

Herrmann, D. N., Horvath, R., Sowden, J. E., Gonzales, M., Sanchez-Mejias, A., Guan, Z., Whittaker, R. G., Almodovar, J. L., Lane, M., Bansagi, B., Pyle, A., Boczonadi, V., Lochmuller, H., Griffin, H., Chinnery, P. F., Lloyd, T. E., Troy Littleton, J., & Zuchner, S. (2014). Synaptotagmin 2 mutations cause an autosomal-dominant form of lambert-eaton myasthenic syndrome and nonprogressive motor neuropathy. American Journal of Human Genetics, 95(3), 332–339. 10.1016/j.ajhg.2014.08.007

Holbrook, S., Finley, J. K., Lyons, E. L., & Herman, T. G. (2012). Loss of syd-1 from R7 Neurons Disrupts Two Distinct Phases of Presynaptic Development. The Journal of Neuroscience, 32(50), 18101. 10.1523/JNEUROSCI.1350-12.2012

Huser, A., Rohwedder, A., Apostolopoulou, A. A., Widmann, A., Pfitzenmaier, J. E., Maiolo, E. M., Selcho, M., Pauls, D., von Essen, A., Gupta, T., Sprecher, S. G., Birman, S., Riemensperger, T., Stocker, R. F., & Thum, A. S. (2012). The Serotonergic Central Nervous System of the Drosophila Larva: Anatomy and Behavioral Function. PLoS ONE, 7(10). 10.1371/journal.pone.0047518

Imlach, W., & McCabe, B. D. (2009). Electrophysiological Methods for Recording Synaptic Potentials from the NMJ of Drosophila Larvae. Journal of Visualized Experiments. 10.3791/1109

Jan, L. Y., & Jan, Y. N. (1976). Properties of the larval neuromuscular junction in Drosophila melanogaster. The Journal of Physiology, 262(1), 189–214. 10.1113/JPHYSIOL.1976.SP011592

Jayakumar, S., Richhariya, S., Venkateswara Reddy, O., Texada, M. J., & Hasan, G. (2016). Drosophila larval to pupal switch under nutrient stress requires IP3R/Ca2+signalling in glutamatergic interneurons. ELife, 5(AUGUST), 1–27. 10.7554/eLife.17495

Jentsch, T. J. (2000). Neuronal KCNQ potassium channels: Physislogy and role in disease. Nature Reviews Neuroscience, 1(1), 21–30. 10.1038/35036198

Kairamkonda, S. (2014). Genetic and molecular characterization of Drosophila melanogaster mutants with compromised motor and reproductive functions (Issue December).

Kairamkonda, S., & Nongthomba, U. (2014). Beadex Function in the Motor Neurons Is Essential for Female Reproduction in Drosophila melanogaster. PLoS ONE, 9(11), e113003. 10.1371/journal.pone.0113003

Kiehn, O., & Kullander, K. (2004). Central Pattern Generators Deciphered by Molecular Genetics. Neuron, 41(3), 317–321. 10.1016/S0896-6273(04)00042-X

Lee, C. W., & Peng, H. B. (2008). The Function of Mitochondria in Presynaptic Development at the Neuromuscular Junction. Molecular Biology of the Cell, 19(January), 150–158. 10.1091/mbc.E07

Li, H., Chaney, S., Forte, M., & Hirsh, J. (2000). Ectopic g-protein expression in dopamine and serotonin neurons blocks cocaine sensitization in Drosophila melanogaster. Current Biology, 10(4), 211–214. 10.1016/S0960-9822(00)00340-7

Lu, Z., Lam, K. S., Wang, N., Xu, X., Cortes, M., & Andersen, B. (2006). LMO4 can interact with Smad proteins and modulate transforming growth factor-β signaling in epithelial cells. Oncogene, 25(20), 2920–2930. 10.1038/sj.onc.1209318

Mahr, A., & Aberle, H. (2006). The expression pattern of the Drosophila vesicular glutamate transporter: A marker protein for motoneurons and glutamatergic centers in the brain. Gene Expression Patterns, 6(3), 299–309. 10.1016/j.modgep.2005.07.006

Marder E., & Bucher D. (2001). Central pattern generators and the control of rhythmic movements. Current Biology, 11(23), R986–R996.

McCabe, B. D., Hom, S., Aberle, H., Fetter, R. D., Marques, G., Haerry, T. E., Wan, H., O’Connor, M. B., Goodman, C. S., & Haghighi, A. P. (2004). Highwire regulates presynaptic BMP signaling essential for synaptic growth. Neuron, 41(6), 891–905. 10.1016/S0896-6273(04)00073-X

Milan, M., Diaz-Benjumea, F. J., & Cohen, S. M. (1998). Beadex encodes an LMO protein that regulates Apterous LIM-homeodomain activity in Drosophila wing development: a model for LMO oncogene function. Genes & Development, 12(18), 2912–2920. 10.1101/gad.12.18.2912

Mohan, J., Firdaus, H., & Nongthomba, UpendraR., R. S. (2015). for Indirect Flight Muscle function . International Journal of Scientific and Research Publications, 5(September).

Moustakas, A., & Heldin, C. H. (2005). Non-Smad TGF-β signals. Journal of Cell Science, 118(16), 3573–3584. 10.1242/jcs.02554

Nahm, M., Lee, M.-J., Parkinson, W., Lee, M., Kim, H., Kim, Y.-J., Kim, S., Cho, Y. S., Min, B.-M., Bae, Y. C., Broadie, K., & Lee, S. (2013). Spartin Regulates Synaptic Growth and Neuronal Survival by Inhibiting BMP-Mediated Microtubule Stabilization. Neuron, 77(4). 10.1016/j.neuron.2012.12.015.Spartin

Nakayama, K., Kiyosue, K., & Taguchi, T. (2005). Diminished neuronal activity increases neuron-neuron connectivity underlying silent synapse formation and the rapid conversion of silent to functional synapses. Journal of Neuroscience, 25(16), 4040–4051. 10.1523/JNEUROSCI.4115-04.2005

O’Connor-Giles, K. M., Ho, L. L., & Ganetzky, B. (2008). Nervous Wreck Interacts with Thickveins and the Endocytic Machinery to Attenuate Retrograde BMP Signaling during Synaptic Growth. Neuron, 58(4), 507–518. 10.1016/j.neuron.2008.03.007

Owald, D., Fouquet, W., Schmidt, M., Wichmann, C., Mertel, S., Depner, H., Christiansen, F., Zube, C., Quentin, C., Körner, J., Urlaub, H., Mechtler, K., & Sigrist, S. J. (2010). A Syd-1 homologue regulates pre- and postsynaptic maturation in Drosophila. Journal of Cell Biology, 188(4), 565–579. 10.1083/jcb.200908055

Patel, M. R., Lehrman, E. K., Poon, V. Y., Crump, J. G., Zhen, M., Bargmann, C. I., & Shen, K. (2006). Hierarchical assembly of presynaptic components in defined C. elegans synapses. Nature Neuroscience, 9(12), 1488–1498. 10.1038/nn1806

Pilling, A. D., Horiuchi, D., Lively, C. M., & Saxton, W. M. (2006). Kinesin-1 and dynein are the primary motors for fast transport of mitochondria in Drosophila motor axons. Molecular Biology of the Cell, 17(4), 2057–2068. 10.1091/mbc.E05-06-0526

Polleux, F., Ince-Dunn, G., & Ghosh, A. (2007). Transcriptional regulation of vertebrate axon guidance and synapse formation. Nature Reviews Neuroscience, 8(5), 331–340. 10.1038/nrn2118

Rawson, J. M., Lee, M., Kennedy, E. L., & Selleck, S. B. (2003). Drosophila neuromuscular synapse assembly and function require the TGF-β type I receptor saxophone and the transcription factor Mad. Journal of Neurobiology, 55(2), 134–150. 10.1002/neu.10189

Rikhy, R., Kumar, V., Mittal, R., & Krishnan, K. S. (2002). Endophilin is critically required for synapse formation and function in Drosophila melanogaster. The Journal of Neuroscience : The Official Journal of the Society for Neuroscience, 22(17), 7478–7484. 10.1523/JNEUROSCI.22-17-07478.2002

Ross, S. E., Greenberg, M. E., & Stiles, C. D. (2003). Basic helix-loop-helix factors in cortical development. Neuron, 39(1), 13–25. 10.1016/S0896-6273(03)00365-9

Salinas, P. C. (2012). Wnt signaling in the vertebrate central nervous system: From axon guidance to synaptic function. Cold Spring Harbor Perspectives in Biology, 4(2), 1–14. 10.1101/cshperspect.a008003

Salinas, P. C., & Zou, Y. (2008). Wnt signaling in neural circuit assembly. Annual Review of Neuroscience, 31, 339–358. 10.1146/annurev.neuro.31.060407.125649

Sandoval, H., Yao, C. K., Chen, K., Jaiswal, M., Donti, T., Lin, Y. Q., Bayat, V., Xiong, B., Zhang, K., David, G., Charng, W. L., Yamamoto, S., Duraine, L., Graham, B. H., & Bellen, H. J. (2014). Mitochondrial fusion but not fission regulates larval growth and synaptic development through steroid hormone production. ELife, 3(October2014), 1–23. 10.7554/eLife.03558

Sanyal, S. (2009). Genomic mapping and expression patterns of C380, OK6 and D42 enhancer trap lines in the larval nervous system of Drosophila. Gene Expression Patterns, 9(5), 371–380. 10.1016/J.GEP.2009.01.002

Shen, K., & Cowan, C. W. (2010). Guidance Molecules in Synapse Formation and Plasticity. Cold Spring Harbor Perspectives in Biology, 1–18.

Shen, W., & Ganetzky, B. (2009). Autophagy promotes synapse development in Drosophila. Journal of Cell Biology, 187(1), 71–79. 10.1083/jcb.200907109

Shoresh, M., Orgad, S., Shmueli, O., Werczberger, R., Gelbaum, D., Abiri, S., & Segal, D. (1998). Overexpression Beadex mutations and loss-of-function heldup-a mutations in Drosophila affect the 3’ regulatory and coding components, respectively, of the Dlmo gene. Genetics, 150(1), 283–299. 10.1093/genetics/150.1.283

Speese, S. D., & Budnik, V. (2012). Sean D. Speese and Vivian Budnik 2007.pdf. 30(6), 268–275. 10.1016/j.tins.2007.04.003.Wnts

Tian, X., Li, J., Valakh, V., Diantonio, A., & Wu, C. (2011). Drosophila Rae1 controls the abundance of the ubiquitin ligase Highwire in post-mitotic neurons. Nature Neuroscience, 14(10), 1267–1275. 10.1038/nn.2922

Trevisan, T., Pendin, D., Montagna, A., Bova, S., Ghelli, A. M., & Daga, A. (2018). Manipulation of Mitochondria Dynamics Reveals Separate Roles for Form and Function in Mitochondria Distribution. Cell Reports, 23(6), 1742–1753. 10.1016/J.CELREP.2018.04.017

Tsai, L. T.-Y., Bainton, R. J., Blau, J., & Heberlein, U. (2004). Lmo mutants reveal a novel role for circadian pacemaker neurons in cocaine-induced behaviors. PLoS Biology, 2(12), e408. 10.1371/journal.pbio.0020408

Ulian-Benitez, S., Bishop, S., Foldi, I., Wentzell, J., Okenwa, C., Forero, M. G., Zhu, B., Moreira, M., Phizacklea, M., McIlroy, G., Li, G., Gay, N. J., & Hidalgo, A. (2017). Kek-6: A truncated-Trk-like receptor for Drosophila neurotrophin 2 regulates structural synaptic plasticity. In PLoS Genetics (Vol. 13, Issue 8). 10.1371/journal.pgen.1006968

Wan, H. I., DiAntonio, A., Fetter, R. D., Bergstrom, K., Strauss, R., & Goodman, C. S. (2000). Highwire regulates synaptic growth in Drosophila. Neuron, 26(2), 313–329. 10.1016/S0896-6273(00)81166-6

Wang, N., Lin, K. K., Lu, Z., Lam, K. S., Newton, R., Xu, X., Yu, Z., Gill, G. N., & Andersen, B. (2007). The LIM-only factor LMO4 regulates expression of the BMP7 gene through an HDAC2-dependent mechanism, and controls cell proliferation and apoptosis of mammary epithelial cells. Oncogene, 26(44), 6431–6441. 10.1038/sj.onc.1210465

Ward, A., Hong, W., Favaloro, V., & Luo, L. (2015). Toll receptors instruct axon and dendrite targeting and participate in synaptic partner matching in a drosophila olfactory circuit. Neuron, 85(5), 1013–1028. 10.1016/j.neuron.2015.02.003

Wentzel, C., Sommer, J., Nair, R., Stiefvater, A., S ibarita, J.-B., & Scheiffele, P. (2013). mSYD1A, a Mammalian Synapse-Defective-1 Protein, Regulates Synaptogenic Signaling and Vesicle Docking. Neuron, 78(6), 1012–1023. 10.1016/j.neuron.2013.05.010.mSYD1A

Williams, C. L., & Smith, S. M. (2018). Calcium Dependence of Spontaneous Neurotransmitter Release. Journal of Neuroscience Research, 96(3), 335. 10.1002/JNR.24116

Williamson, W. R., & Hiesinger, P. R. (2008). Neuroscience: Synaptic patterning by morphogen signaling. Science Signaling, 1(18), 18–21. 10.1126/stke.118pe20

Wu, C., Wairkar, Y. P., Collins, C. A., & DiAntonio, A. (2005). Highwire function at the Drosophila neuromuscular junction: Spatial, structural, and temporal requirements. Journal of Neuroscience, 25(42), 9557–9566. 10.1523/JNEUROSCI.2532-05.2005

Zeng, C., Justice, N. J., Abdelilah, S., Chan, Y.-M., Jan, L. Y., & Jan, Y. N. (1998). The Drosophila LIM-only gene, dLMO, is mutated in Beadex alleles and might represent an evolutionarily. Proceedings of the National Academy of Sciences of the United States of America, 95(September), 10637–10642.

Zenvirt, S., Nevo-Caspi, Y., Rencus-Lazar, S., & Segal, D. (2008). Drosophila LIM-Only Is a Positive Regulator of Transcription During Thoracic Bristle Development. Genetics, 179, 1989–1999. 10.1534/genetics.108.090076

