## Supplemental Figures for "Beadex, the *Drosophila* LIM only protein, is required for the growth of the larval neuromuscular junction and sensorimotor activities"

SUPPLEMENTARY FIGURE 1

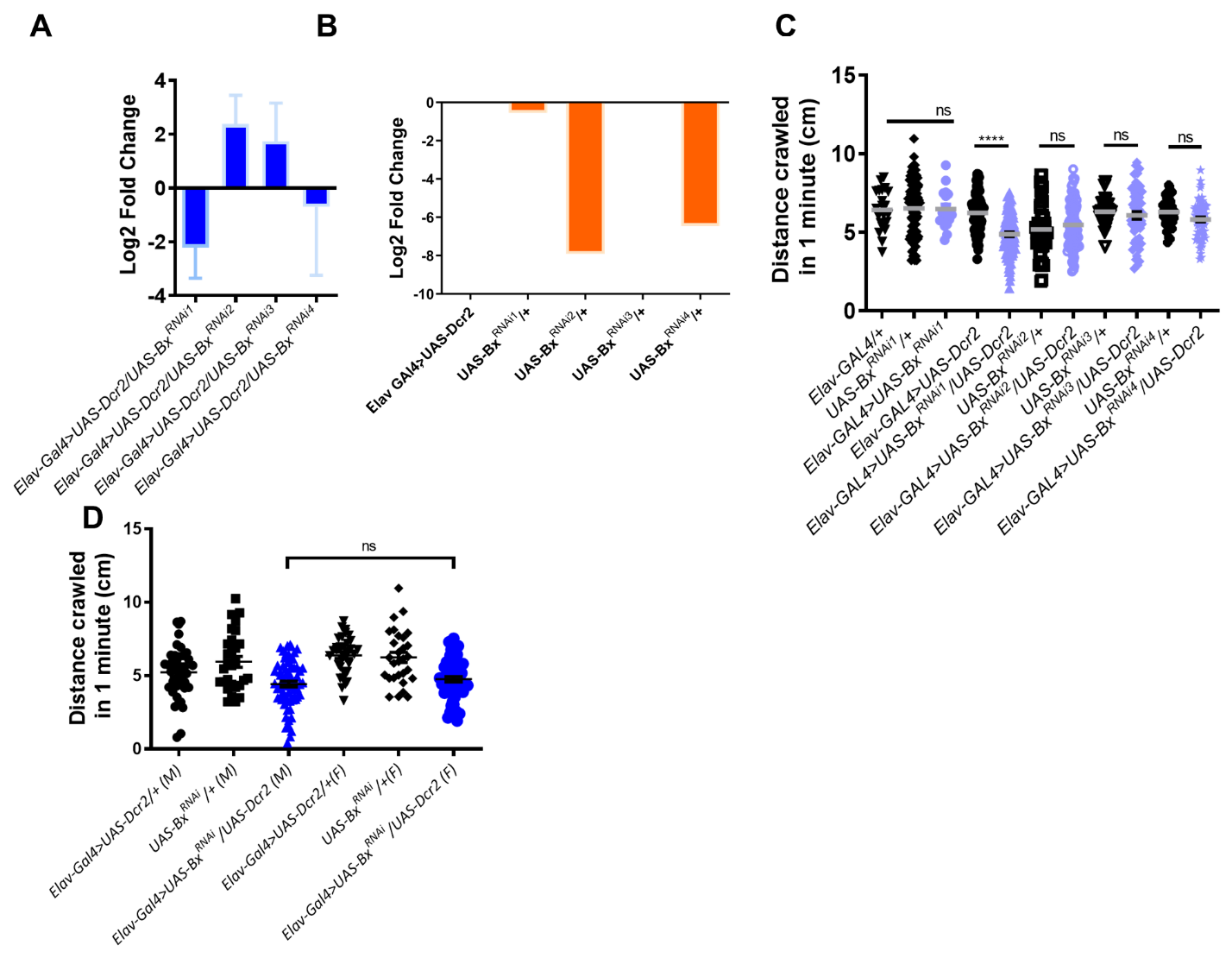

SUPPLEMENTARY FIGURE 2

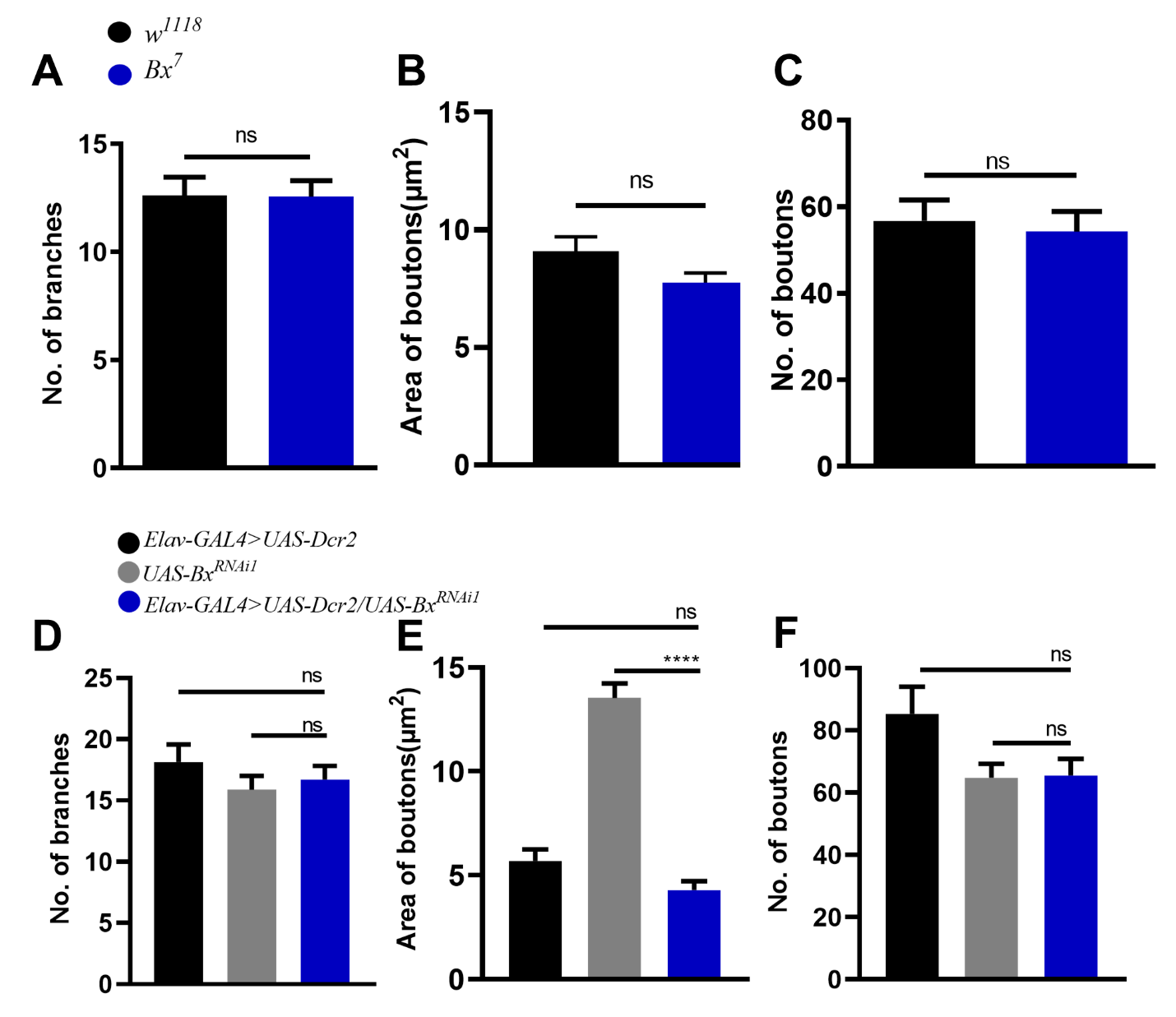

SUPPLEMENTARY FIGURE 3

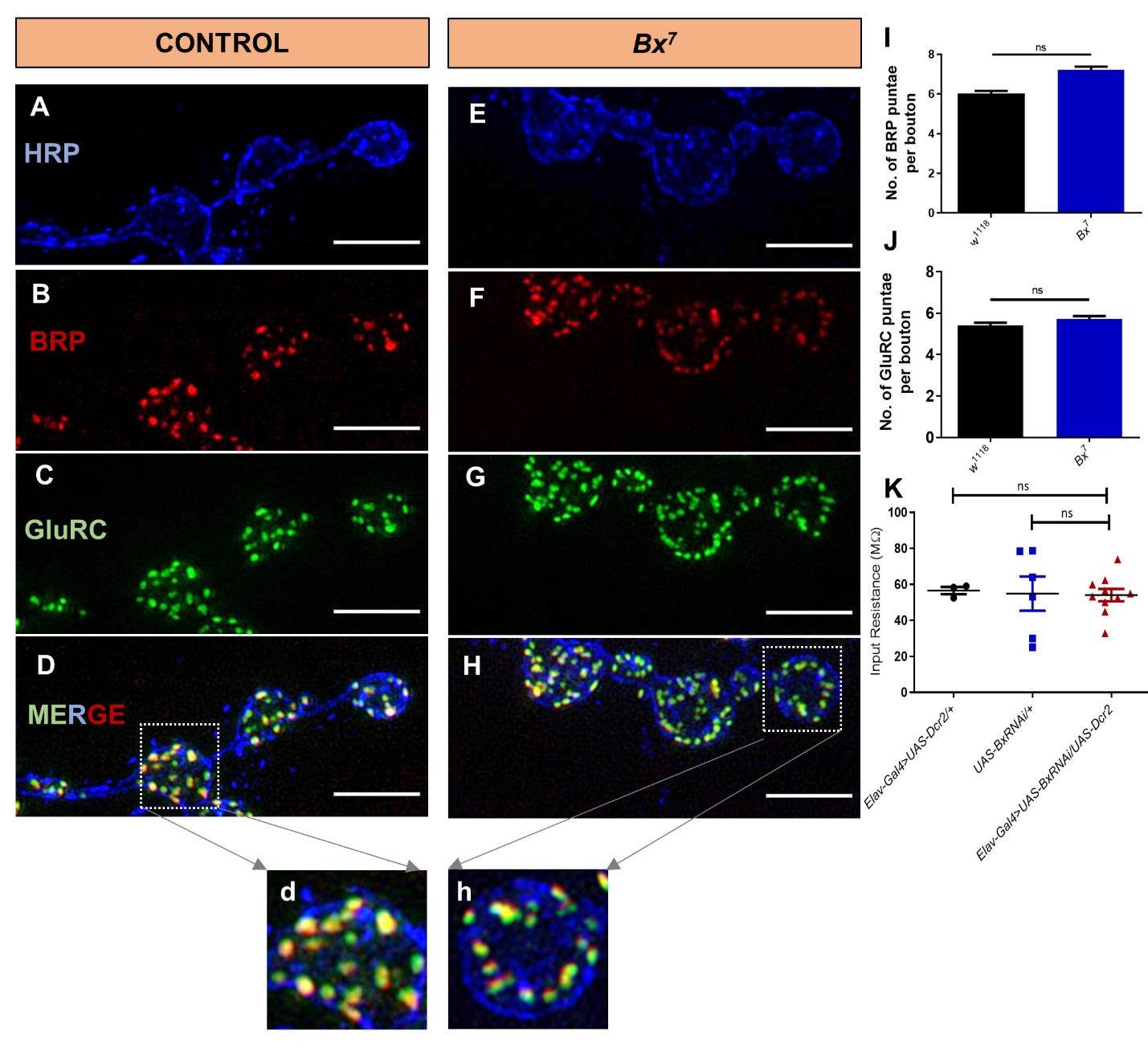

SUPPLEMENTARY FIGURE 4

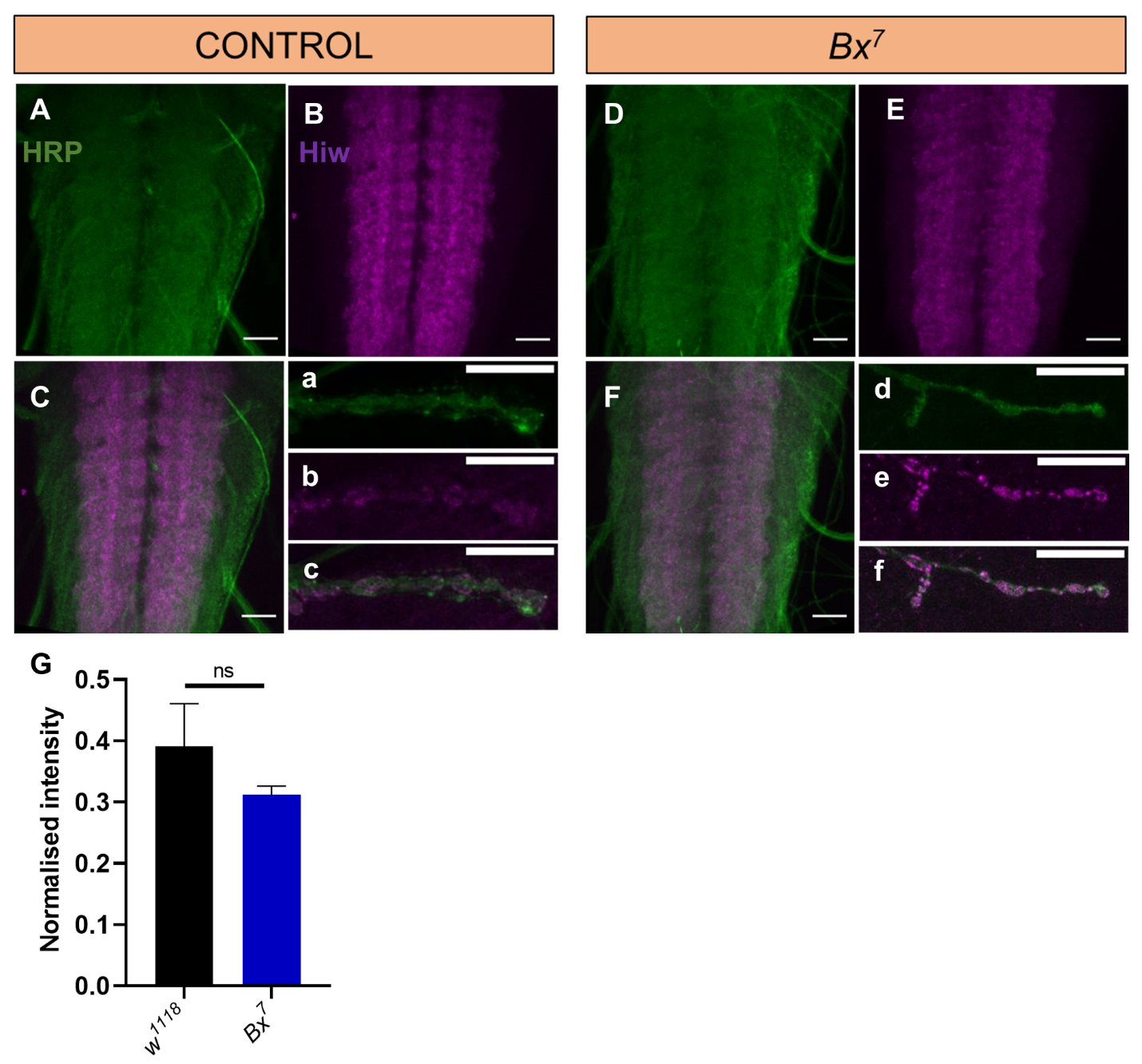

SUPPLEMENTARY FIGURE 5
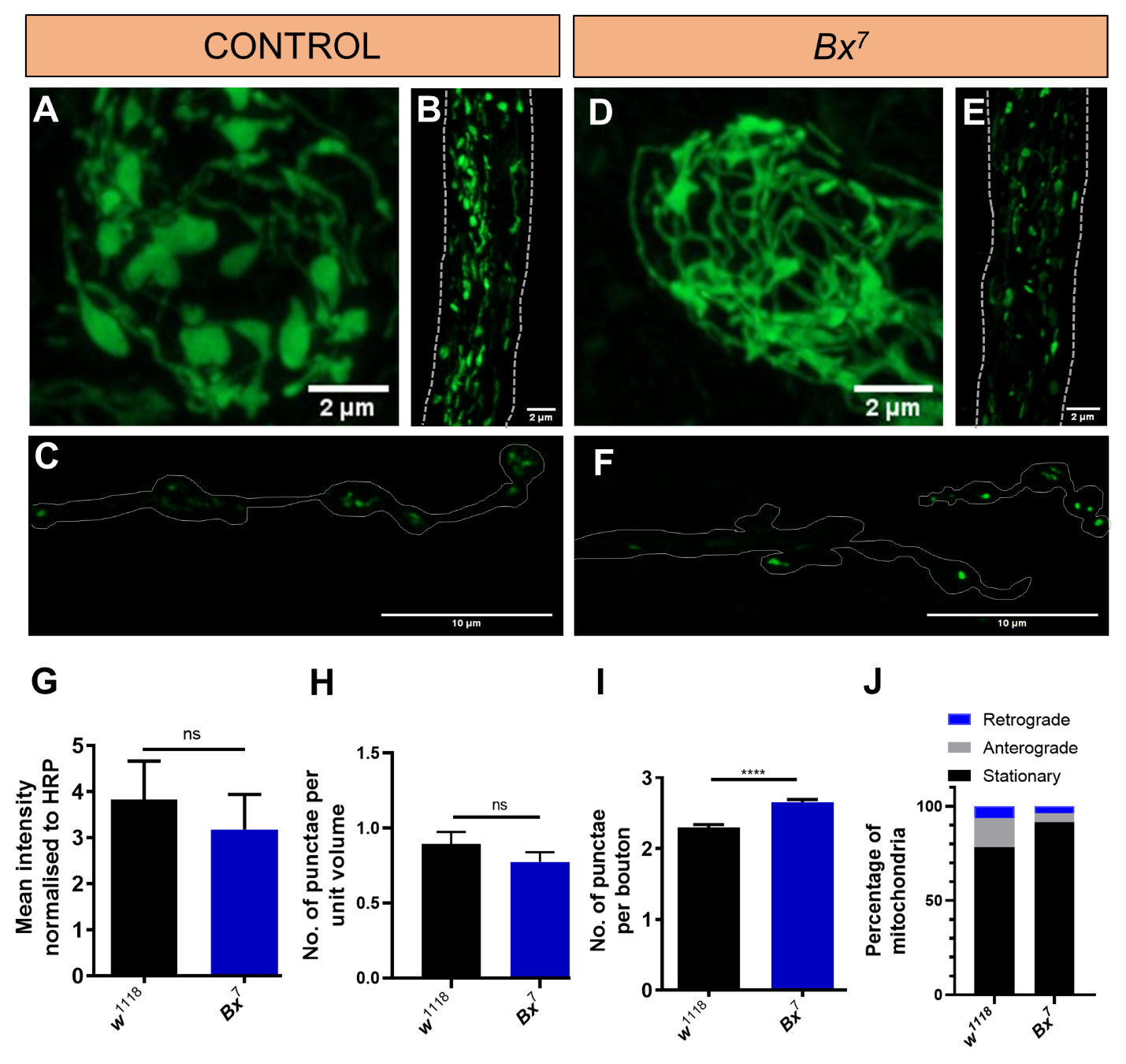
