## Supplementary figures and images for "Beadex, the *Drosophila* LIM only protein, is required for the growth of the larval neuromuscular junction and sensorimotor activities"

### Supplemental Tables

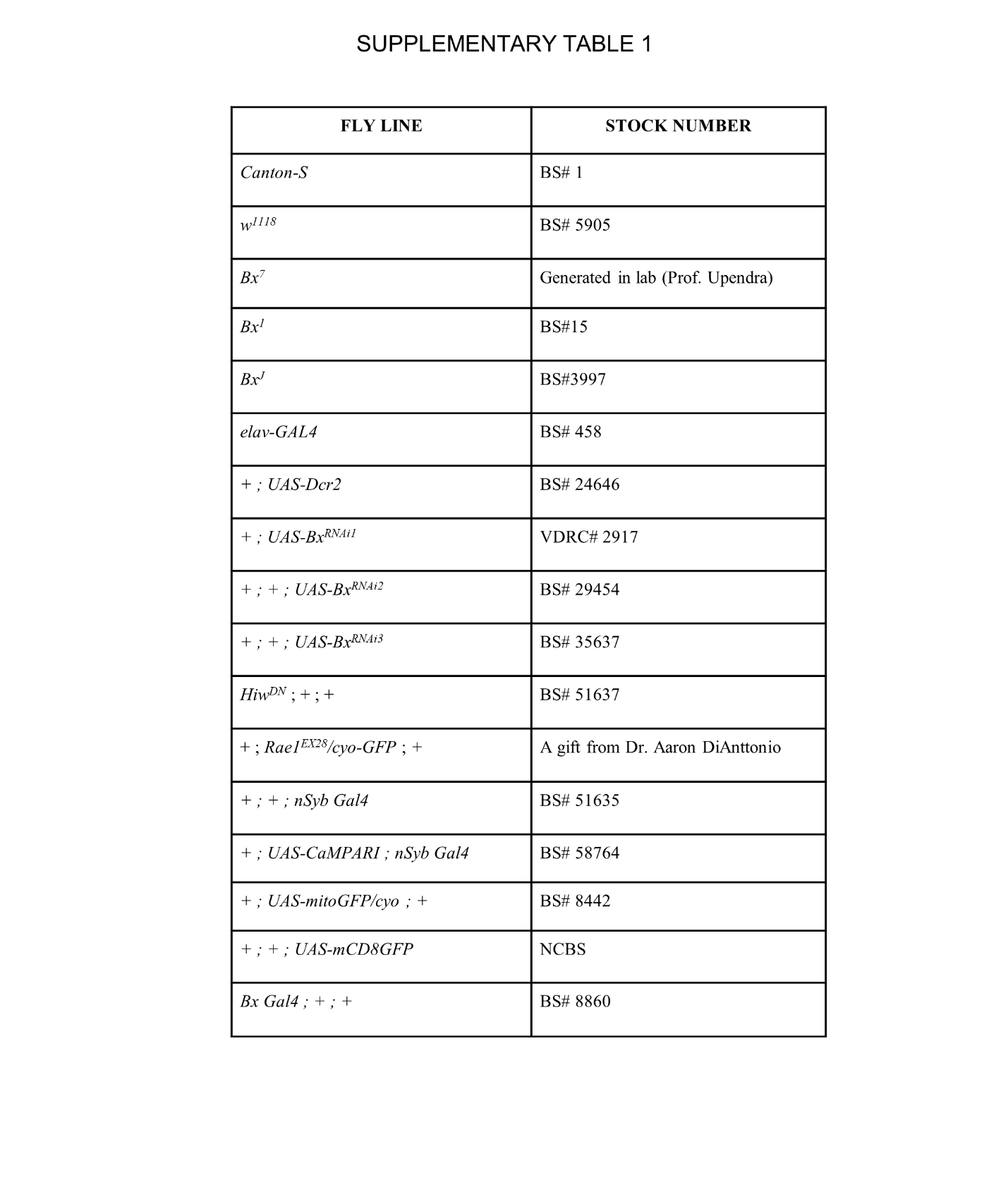

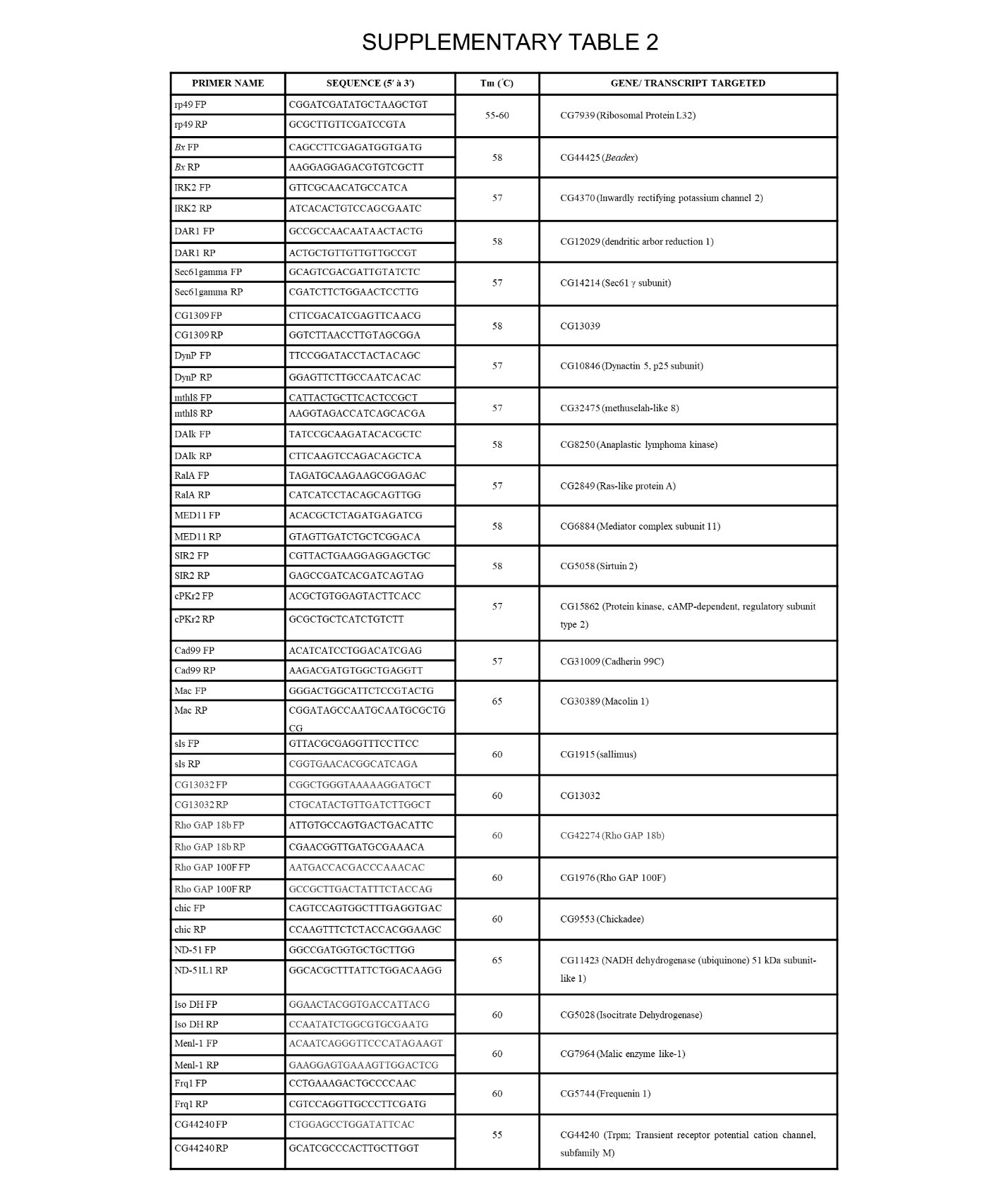
